# Amphiphile-engineered DNA adjuvants stimulate strong type I IFN production in lymph nodes via cytosolic danger-sensing to induce potent cellular and humoral immunity in mice and non-human primates

**DOI:** 10.1101/2024.11.01.621501

**Authors:** Martin P. Steinbuck, Lochana M. Seenappa, Wei Zhan, Erica Palmer, Aniela Jakubowski, Xavier Cabana-Puig, Mimi M. Jung, Lisa K. McNeil, Christopher M. Haqq, Katherine A. Fitzgerald, Peter C. DeMuth

## Abstract

Adjuvants are immuno-activators capable of shaping the magnitude and quality of antigen-specific immune responses induced by subunit immunization. Presently, there is an acute need for effective adjuvants that safely induce durable and balanced humoral and cellular responses; the latter being indispensable for protection against intracellular pathogens and cancer. Here, we iteratively optimized a novel class of Amphiphile (AMP)-modified, immunostimulatory DNA-adjuvants designed for targeted delivery to lymph nodes and enhanced stimulation of cytosolic danger-sensing pathways to generate strong adaptive immunity. AMP-DNA adjuvants induced potent IFN-I-driven inflammatory environments in mouse and NHP lymph nodes that were dependent on TBK1 signaling, leading to significantly enhanced cytokine secretion by polyfunctional CD8^+^ and CD4^+^ T cells in multiple tissues, and strongly elevated T_H_1-associated and neutralizing antibody responses, without toxicity. These results demonstrate that AMP-engineering enables lymph node-targeted DNA-adjuvants to uniquely activate cytosolic immune-signaling to generate robust adaptive responses crucial for vaccine efficacy.

## INTRODUCTION

Vaccines have transformed public health since their inception and remain one of the most cost-effective and widely applied health interventions available. At present, modern protein subunit vaccines utilize highly defined antigens to elicit precise and safe immune responses against their specific targets. However, such refined antigens, in isolation, often lack ancillary immunostimulatory elements that engage the innate immune system to induce robust and lasting immunity. Thus, besides a few exceptions^1,2^, adjuvants have inevitably become indispensable components of subunit vaccines essential for stimulating adequate responses to target antigens^3^. Whereas some adjuvants mainly operate as antigen delivery or depot systems, others work as immunostimulants that activate antigen presenting cells (APC) through toll-like receptors (TLRs) or other pattern recognition receptors (PRRs)^4^. To date, the most commonly used adjuvants in commercially available prophylactic vaccines primarily function as antigen depots which predominantly amplify antibody responses (e.g. aluminium salts, oil-in-water emulsions)^5^. Notable disadvantages of these classical adjuvants often include the lack of long-term protective immunity^4,6^ and a poor ability to induce cellular immune responses^7^ critical for protection against intracellular pathogens such as malaria^8,9^, tuberculosis^10,11^, and viruses, as well as cancer^12^.

In contrast, many molecular adjuvants, including TLR-agonists, are promising adjuvant candidates as they often stimulate type I interferon (IFN-I) and inflammatory cytokine responses, which have the potential to induce strong T_H_1-associated cellular immunity^4^. Nonetheless, only a small number of these molecules have been tested in clinical trials and approved for use in humans, including synthetic double stranded RNA (e.g. poly(I:C)/poly-ICLC, a TLR3-agonist), oligodeoxynucleotides (ODN; e.g. CpG-7909, a TLR9-agonist), cyclic dinucleotides (e.g. cyclic GMP-AMP, a STING-agonist), and nucleoside analogs (e.g. resiquimod, a TLR7/8-agonist). A common obstacle with molecular adjuvants is their tendency to either remain in the injection site (molecules >100 kDa) or to be distributed systemically (<20 kDa), as low-molecular weight compounds readily diffuse into circulation through capillary endothelial tight junctions^13^. This often leads to rapid systemic dissemination towards potentially immunologically irrelevant or tolerizing sites. Systemic exposure of adjuvants thus makes it difficult to achieve sufficient immunostimulation while avoiding reactogenicity, necessitating dose restrictions which limit their use in practice^14–16^.

However, recent innovations in adjuvant design and the use of delivery platforms including engineered nanoparticles have contributed to increased adjuvant activity, vaccine immunogenicity, and improved safety^17,18^. Among these efforts are technologies that aim to enhance the biodistribution and thus potency of vaccines by targeting delivery directly to lymph nodes (LN), where protective immune responses are coordinated^19^. One such approach is “albumin-hitchhiking”, which utilizes albumin’s size-dependent (∼65 kDa) migration through the lymphatics, to accumulate vaccines in draining LNs^20,21^. The resulting targeted bioavailability and increased APC uptake of antigens and adjuvants consequently initiates robust adaptive immune responses. Accordingly, to allow for transient association with albumin, antigens and adjuvants can be chemically modified with albumin-binding moieties^22–24^. We have particularly focused on the preclinical and clinical development of amphiphilic lipid polymer (Amphiphile; AMP)-conjugates, wherein a molecular payload of interest is linked to an albumin-binding phospholipid tail, enabling noncovalent association with endogenous albumin following injection into tissue that mediates efficient targeting to draining LNs via the lymphatics. AMP-conjugates have previously been demonstrated in mice and non-human primates (NHP) to optimize *in vivo* biodistribution and prevent systemic toxicity of the TLR9-agonist CpG-7909 in the context of SARS-CoV-2 vaccination, while generating strong, long-term cellular and humoral immunity^25,26^. Furthermore, ELI-002, an AMP-CpG adjuvanted peptide vaccine targeting mutant KRAS antigens that drive colorectal and pancreatic cancers was recently evaluated in a phase 1 clinical trial (AMPLIFY-201; NCT04853017) and an ongoing phase 1/2 trial (AMPLIFY-7P; NCT05726864), showing that AMP-conjugated adjuvants are well tolerated and induce robust anti-tumor T cell responses associated with tumor biomarker reduction that correlate with prolonged relapse-free survival in humans^27^.

To build on the potential of this approach, we sought to develop additional adjuvants that utilize signaling pathways distinct from CpG:TLR9. Cytosolic DNA is a promising candidate because of its strong induction of inflammation if detected outside of the nucleus or mitochondria. The cytosol contains numerous nucleic acid sensing PRRs^28^, that can detect a wide range of foreign and host DNA, generating robust responses by the innate immune system. Nonetheless, to utilize DNA as a vaccine adjuvant there are many drug delivery challenges that must be overcome, such as the negligible cellular uptake, rapid clearance, and poor cytosolic access of freely administered DNA^29^, which may explain why the development of these DNA-based intracellular PRR stimulators has been limited thus far. Therefore, targeted delivery of DNA adjuvants to the appropriate innate immune cells *in vivo* combined with efficient cytosolic entry that allows pharmacological activation of cognate DNA sensors will provide new opportunities with significant potential to elicit strong immune responses.

In this study, we designed a series of AMP-engineered DNA adjuvants capable of inducing highly potent CD8^+^ and CD4^+^ T cell immunity against co-administered protein subunit antigens in mice and NHPs. AMP-induced cellular immune responses significantly exceeded those of comparator vaccines adjuvanted with unmodified ‘soluble’ (SOL) DNA or commercially available adjuvants. Moreover, AMP-mediated production of substantial neutralizing IgG1 titers was accompanied by increased levels of T_H_1-associated IgG2c, an isotype known to facilitate cellular immunity. This was preceded by strong induction of innate responses generating a highly inflammatory, LN-confined milieu resulting in enhanced PRR-sensing, and antigen-processing and -presentation capabilities. These mechanisms were critically reliant on IFN-I, which in turn depended on TANK-binding kinase 1 (TBK1) signaling, as demonstrated by IFNα receptor-1 (IFNAR1)-blockade and studies in mice lacking TBK1 expression in hematopoietic cells. Overall, this study demonstrates the potency of AMP-DNA adjuvants to stimulate vigorous cellular and humoral immune responses and emphasizes the versatility of AMP-modification to efficiently deliver otherwise ineffective adjuvants to potentiate their immunogenicity.

## RESULTS

### AMP-DNA is a potent and safe adjuvant inducing T cell and Ab responses to protein subunit antigens in mice

DNA-adjuvants were designed with the objective to stimulate innate immune activation by engaging intracellular nucleic acid sensing PRRs. Multiple design parameters were assessed, including DNA 5’ AMP-modification, DNA-strandedness, oligonucleotide sequence, length, and backbone-linkage chemistry. To assess the importance of AMP-modification on the immunogenicity of DNA adjuvants, we first synthesized phosphorothioate-backbone, single- or double-stranded AMP-DNA, consisting of a 5’ diacyl phospholipid-conjugated, single-stranded 50-mer polythymidine ODN alone (AMP-dT_50_) or annealed to a complementary, unconjugated polyadenosine strand (AMP-dT_50_ + dA_50_ → AMP-dA:dT_50_; Fig. 1a; S1a). These were compared to their corresponding unmodified SOL DNA analogues. The DNA adjuvants were admixed (Fig. 1b) with either ovalbumin (OVA), a research antigen commonly used to benchmark relative vaccine immunogenicity, or SARS-CoV-2 WH-01-variant spike protein receptor-binding domain (WH-01 RBD), representing a highly relevant antigen of public concern. Immunization was conducted in a biweekly prime-boost regimen on day 0 and day 14, followed 7 days later by analysis of cellular and humoral immune responses (Fig. 1c; S2a). Immunization with AMP-DNA and OVA resulted in a 7- and 41-fold increase in the mean number of splenic IFNγ spot-forming cells (SFC) for AMP-dA:dT_50_ (3,962 SFC/1×10^6^) and AMP-dT_50_ (6,934 SFC/1×10^6^), respectively, compared to their unmodified counterparts which showed no signal above mock-treated mice (Fig. 1d). Moreover, these responses were polyfunctional, exhibiting significantly increased antigen-specific secretion of IFNγ (198-fold), TNFα (19-fold), GM-CSF (108-fold), IL2 (5-fold), and granzyme B (GzmB, 44-fold), when compared to SOL vaccine, which induced responses equivalent to mock treatment (Fig. 1e,f). Further, tetramer and intracellular cytokine staining of peripheral blood mononuclear cells (PBMC) showed that AMP-dA:dT_50_ or AMP-dT_50_ immunization elicited substantial circulating CD8^+^ T cell responses recognizing the immunodominant OVA-epitope, SIINFEKL (65% and 80% of circulating CD8^+^ T cells, respectively; Fig. 1g) and that CD8^+^ and CD4^+^ T cells were key contributors of cytokine secretion, with 62% and 77% of CD8^+^ T cells actively producing cytokines (Fig. 1h,i; S1b). Similar outcomes were observed in T cells isolated from perfused lung tissue (Fig. S1c,d), an organ that is a frequent point of entry for pathogens and a primary organ for tumor metastasis (gating strategies are depicted in Fig S1e,f). In addition to strong cellular responses, AMP-modified adjuvants generated high serum IgG titers, while SOL groups produced lower, but nonetheless robust OVA-specific antibody levels (Fig. 1j). This may suggest that while humoral responses can be achieved with soluble, unmodified DNA adjuvants, AMP-DNA adjuvants provide specific benefits to overcome the more rigorous threshold of activation for robust cellular responses. Analogous results were observed against the RBD antigen (Fig. S2), where splenic IFNγ secretion was enhanced by AMP-dA:dT_50_ and AMP-dT_50_ over their SOL DNA adjuvant comparators by 7- and 12-fold, respectively (Fig. S2b) and resulted in antigen-specific cytokine production in 65% and 85% of circulating CD8^+^ T cells (Fig. S2d); soluble comparators were inactive. Furthermore, analysis of long-term cellular immunity after AMP-DNA immunization in both RBD and OVA models demonstrated durable antigen-specific CD8^+^ T cell responses and an expanding population of central memory T cells over time (Fig. S3). No adverse body weight changes or systemic cytokine levels were observed upon immunization with AMP-DNA, and the primary cytokines implicated in cytokine release syndrome, namely IL1/2/6/10, IFNγ, GM-CSF and TNFα, were not detected at high levels in the serum (Fig. S4). Together, these data illustrate the potency of AMP-DNA as a powerful and safe vaccine adjuvant that can generate strong, durable cellular and humoral immune responses.

**Figure 1:**
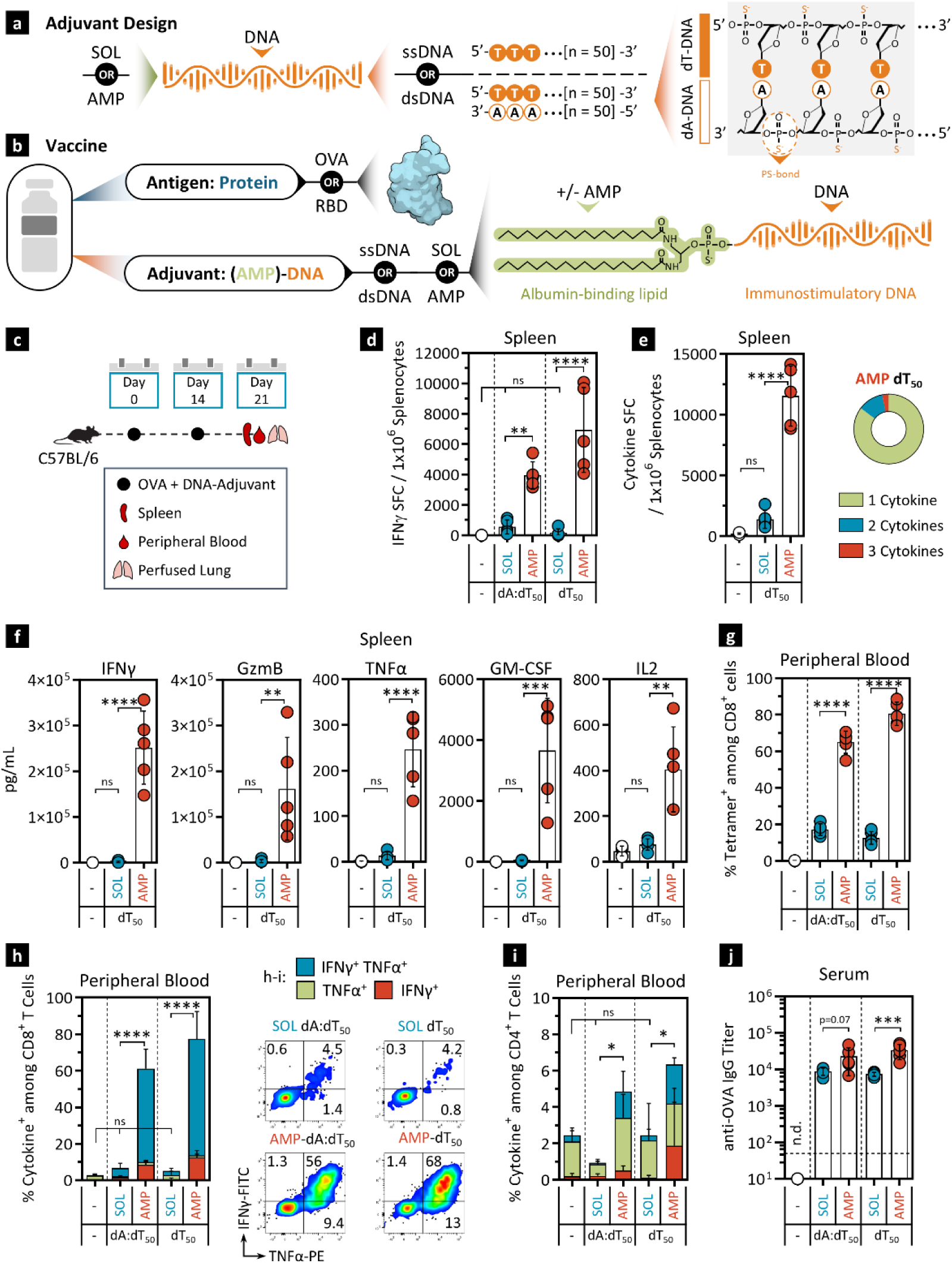
AMP-DNA is a potent adjuvant for inducing T cell and Ab responses to protein subunit antigens. **a)** Schematic drawings depicting the chemical structure of the AMP-DNA adjuvants. The ssDNA variant, dT_50_ consists of 50 phosphorothioate (PS)-linked deoxythymidines. When annealed to a 50-mer deoxyadenosine DNA the resulting dsDNA forms dA:dT_50_. The diacyl lipid (AMP-tail) is conjugated to the 5’-end of dT_50_ in both ssDNA and dsDNA forms. **b)** Vaccine formulation consists of the protein antigen (OVA or RBD) admixed with the candidate DNA-adjuvants depicted in a) and Fig. S1a), and ISD or HSV_60_. **c)** Schema showing animal dosing and experimental schedule. C57BL/6J mice (n = 5) were immunized twice with 5 μg OVA protein and 5 nmol SOL or AMP-DNA adjuvants. Cells were assayed 7 days post booster dose. **d)** ELISpot analysis of splenocytes restimulated with OVA OLPs overnight and assayed for IFNγ production. Shown is the frequency of IFNγ SFCs per 10^6^ splenocytes. **e)** FluoroSpot analysis of splenocytes restimulated with OVA OLPs overnight and assayed for IFNγ, TNFα and IL2 production. Shown is the frequency of SFCs per 10^6^ splenocytes that produce at least one cytokine. Donut charts represent percentages of polyfunctional versus single cytokine secreting cells. **f)** Multiplexed proteomics analysis (Luminex) of splenocytes restimulated with OVA OLPs overnight and assayed for IFNγ, TNFα, IL2, GM-CSF and GzmB production. Depicted are quantities of secreted cytokines in the culture supernatant. **g)** Peripheral blood CD8^+^ T cells were assayed *ex vivo* for OVA reactive TCRs using a SIINFEKL-specific tetramer. **h-i)** Flow cytometric analysis of cytokine production by CD8^+^ (g) and CD4^+^ (h) T cells in peripheral blood. Shown are percentages of cytokine^+^ cells among CD8^+^ or CD4^+^ T cells and representative flow cytometry scatter plots of IFNγ and TNFα positive CD8^+^ T cells. **j)** Serum IgG titers were determined against OVA protein. Mock vaccines contained vehicle only. Values depicted are means±SD. *p < 0.05, **p < 0.01, ***p < 0.001, ****p < 0.0001 by one-way ANOVA followed by Tukey’s post-hoc analysis. n.d., not detected

### Immunogenicity of AMP-DNA adjuvants depends on key physical and chemical properties

To establish additional physical and chemical characteristics beyond AMP-modification that determine optimal immunogenicity of AMP-DNA adjuvants, the remaining four initially selected design parameters (Fig. 2a) were systematically altered to determine their contribution to the resulting immune response. Immunogenicity was determined by measuring antigen-specific IFNγ secretion in splenic T cells. Examining DNA-strandedness demonstrated that both single-stranded and double-stranded DNA induced robust cytokine production in antigen-specific T cells. While single-stranded AMP-dT generated stronger immune responses than its double-stranded form, AMP-dA:dT (Fig. 2b), two other well-established immunostimulatory ODNs^30^, herpes simplex virus 60 (HSV_60_) and *Listeria monocytogenes*-derived IFN-stimulatory DNA (ISD) performed equally well in either configuration upon AMP-conjugation, thus substantiating that strandedness may not be a crucial characteristic for adjuvant recognition (Fig. 2c,d). These results also suggest that cellular DNA-sensing may be largely sequence-agnostic as significant responses were observed to all aforementioned sequences. In contrast, single stranded polyadenosine (AMP-dA) as well as AMP-dT:dA, which contains the AMP-modification on the dA strand, were unable to generate robust immunity (Fig. 2b). However, an ODN comprised of alternating thymidine and adenosine bases (dAdT:dAdT) was able to substantially rescue immune responses (Fig. S5), suggesting that certain sequence requirements do exist and that thymidine containing sequences may be preferred by the relevant DNA sensing PRRs. Another crucial factor known to determine DNA detection by PRRs is ODN length^30^. Systematic variation of sequence length for AMP- and SOL-DNA showed that sequences shorter than 30 nucleotides were unable to generate T cell responses (Fig. 2e,f), indicating that the involved PRRs require a minimum sequence length to transmit the danger signal. However, increasing DNA length beyond 30-50 nucleotides did not confer additional benefits. Lastly, exchanging the synthetic phosphorothioate (PS) bonds with naturally occurring phosphodiester (PO) bonds abrogated all cellular immune responses (Fig. 2g,h), possibly due to poor stability *in vivo* of PO-bonds due to nuclease-dependent degradation. Overall, these data suggest that the optimal single- or double-stranded AMP-DNA adjuvant should contain 30-50 PS-linked nucleotides of which a sufficient number are thymidine for effective PRR recognition.

**Figure 2:**
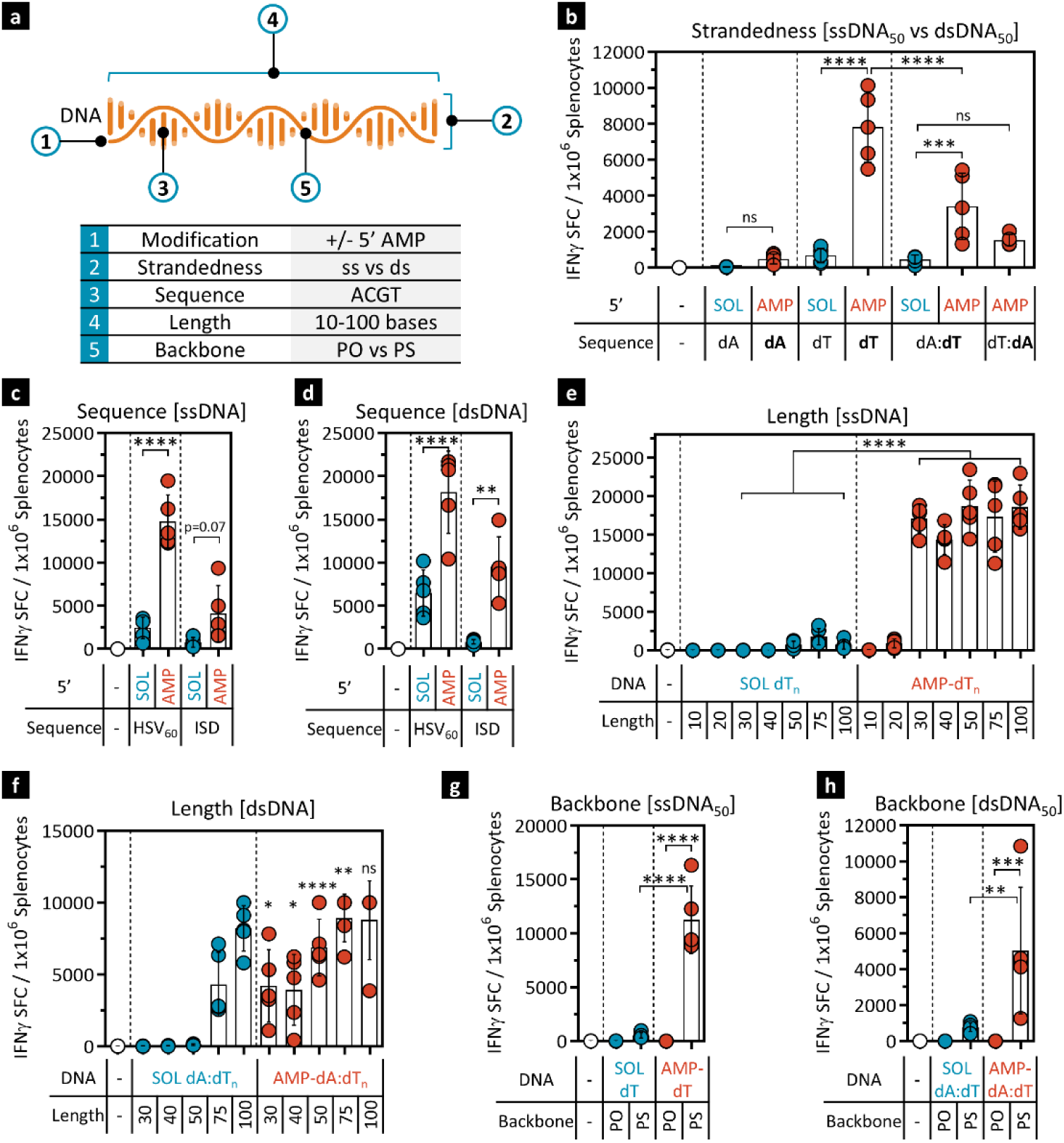
DNA design features dictate the strength of resulting immunogenicity. C57BL/6J mice (n = 5) were immunized as described in Fig 1c. Immunizations contained 5 μg WH-01 RBD protein and 5 nmol SOL or AMP-DNA adjuvant. Cells were assayed 7 days post booster dose. **a)** Schematic describing the modifications of the DNA adjuvants. **b-h)** Evaluation of the effect of DNA modifications on immune activation. Splenocytes were restimulated with RBD OLPs overnight and assayed for IFNγ production by ELISpot. Shown is the frequency of IFNγ SFCs per 10^6^ splenocytes; The assessed DNA modifications included: b) single stranded versus double stranded DNA, (bold letters indicate the AMP-conjugated strand); c-d) HSV_60_ and ISD sequences in single stranded (c) and double stranded (d) form, e-f) DNA length variants for dT (e) and dAdT (f); g-h) phosphodiester (PO)- vs phosphorothioate (PS)-linked bases for dT (g) and dAdT (h). Mock vaccines contained vehicle only. Values depicted are means±SD. *p < 0.05, **p < 0.01, ***p < 0.001, ****p < 0.0001 by one-way ANOVA followed by Tukey’s post-hoc analysis. Statistical analyses in panels e and f compare each AMP-modified length variant to its soluble counterpart.

To further elucidate the importance of adjuvant design on immunogenicity, *in vivo* imaging was conducted in which DNA was 3’-conjugated to a Cy5 fluorescent tag to allow analysis of adjuvant biodistribution upon injection. dT variants of 50 and 20 nucleotides in length were selected, representing examples with potent immunogenic activity or complete lack thereof, respectively. As expected, given their hydrodynamic size (>10nm), both SOL and AMP-dT_50_ efficiently accumulated in the inguinal LNs^13^, whereas the shorter SOL dT_20_ failed to do so (Fig. S6a-c). AMP-modification, however, was able to reinstate efficient delivery of AMP-dT_20_, consistent with prior observations for similarly sized CpG-7909^22^. This suggests that DNA molecules of small size are poor adjuvants partly due to their untargeted systemic distribution, with only small quantities reaching immunologically relevant cells within the LNs. In addition, cytosolic DNA-detection requires sufficient sequence length, as even when AMP-dT_20_ reaches the LN, it is incapable of stimulating an immune response (Fig. 2c). Notably, SOL dT_50_ as well as SOL dA:dT_50_ (Fig. S6d,e) were incapable of eliciting responses despite reaching the LNs and being of sufficient length. Therefore AMP-modification may facilitate additional mechanisms beyond LN delivery which are critical for immune activation: improved cellular uptake by APCs, endocytic escape leading to enhanced cytosolic PRR access, or other mechanisms promoting cognate PRR engagement are possible factors of great interest for future studies.

### AMP-DNA induces superior cellular and humoral immunogenicity relative to clinical benchmark adjuvants

To test how AMP-DNA adjuvants compare to commercially available and/or clinically used adjuvants, a panel was selected that included examples of common vaccine adjuvants available for use in humans. These included Alhydrogel (aluminium hydroxide gel; alum); a group of oil-and-water emulsions including incomplete Freund’s adjuvant (IFA), AddaVax (an MF59 mimetic), and AddaS03 (an AS03 mimetic); as well as the TLR4 agonist monophosphoryl Lipid A (MPLA, an LPS derivative)^4,31^. AMP-CpG-7909 was also included in this panel to compare among AMP-conjugated adjuvants. AMP-DNA consistently produced significantly higher cellular responses than any of the benchmark adjuvants or even AMP-CpG-7909. Both AMP-dT_50_ and AMP-dA:dT_50_ generated greater numbers of circulating antigen-specific CD8^+^ T cells (Fig. 3a) and increased frequencies of cytokine^+^ CD8^+^ and CD4^+^ T cells in peripheral blood, splenic and perfused lung samples (Fig. 3b-f), whereas comparator immunization did not elicit any notable cellular immunity. In contrast, all adjuvants were able to generate high titers of anti-OVA serum IgG (Fig. 3g). However, more detailed analysis of specific IgG subclasses, namely IgG_1_ and IgG_2c_ (an IgG_2a_ paralog in C57BL/6 mice^32^), showed differences in isotype profiles. While titers of IgG_1_, whose main function is to neutralize soluble antigens, were comparable in all adjuvant groups tested, AMP-DNA and AMP-CpG vaccines generated 1-2 orders of magnitude higher levels of IgG_2c_, an IFNγ-induced isotype that strongly promotes complement-fixing and antibody-dependent cellular cytotoxicity^33^ (Fig. 3h-j). This is likely a result of the significant levels of IFNγ produced by effector T cells in these mice. Overall, while most well-established adjuvants were limited to T_H_2-associated humoral responses, AMP-conjugated DNA-adjuvants were capable of complementing robust humoral with strong cellular immune activation.

**Figure 3:**
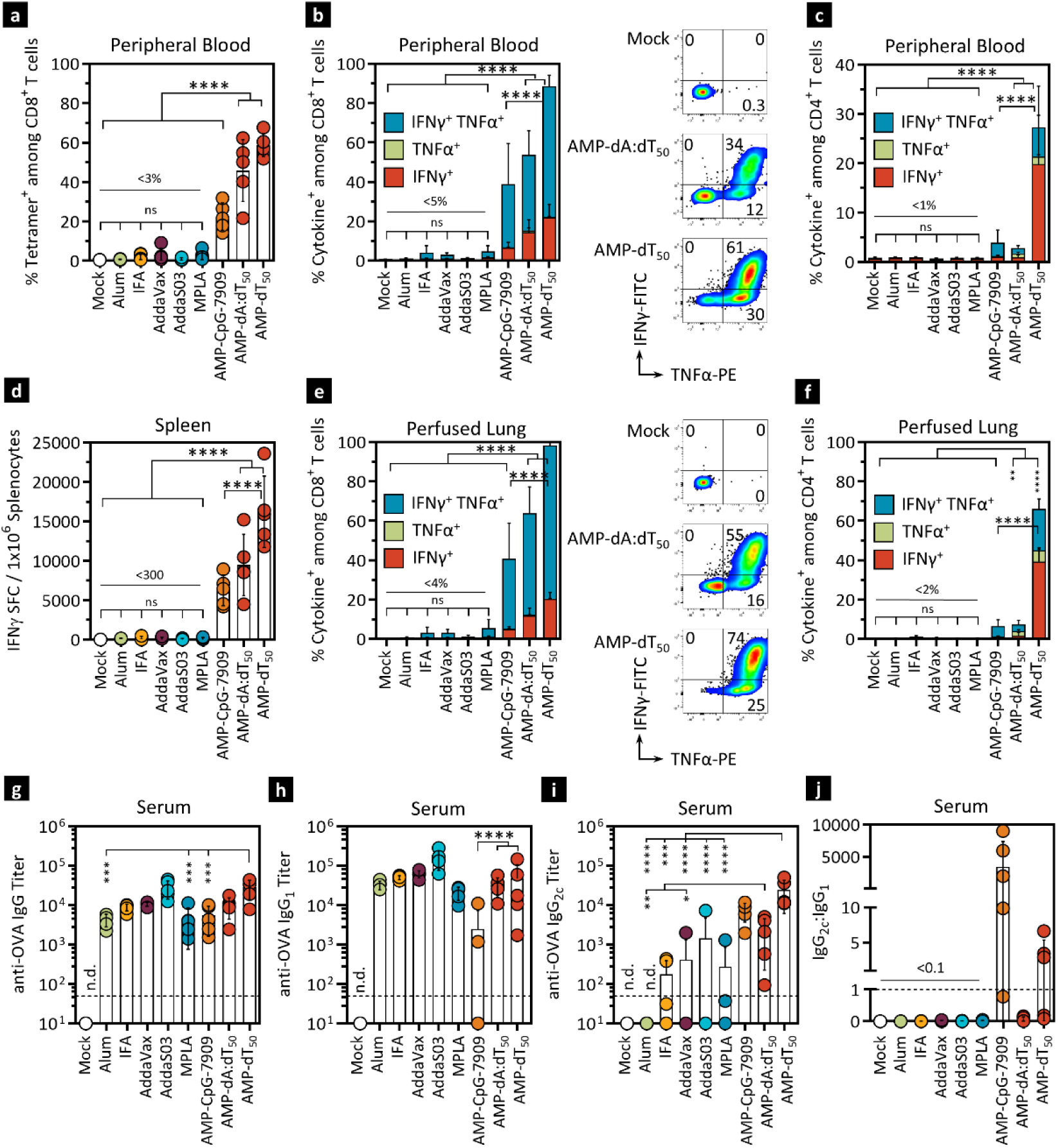
AMP-DNA induces superior cellular and humoral immunogenicity relative to clinical benchmark adjuvants. C57BL/6J mice (n = 5) were immunized as described in Fig 1c. Immunizations contained 5 μg OVA protein and either 5 nmol AMP-DNA, 1 nmol AMP-CpG-7909, or the indicated comparator adjuvants (100 μg alum; 10 μg MPLA; 1:2 emulsions of IFA, AddaVax, AddaS03). Cells were assayed 7 days post booster dose. **a)** Peripheral blood CD8^+^ T cells were assayed *ex vivo* for OVA-reactive TCRs using a SIINFEKL-specific tetramer. **b-c)** Flow cytometric analysis of cytokine production by CD8^+^ (b) and CD4^+^ T cells (c) in peripheral blood. Shown are percentages of cytokine^+^ cells among CD8^+^ or CD4^+^ T cells and representative flow cytometry dot plots of TNFα and IFNγ positive CD8^+^ T cells. **d)** ELISpot analysis of splenocytes restimulated with OVA OLPs overnight. Shown is the frequency of IFNγ SFCs per 10^6^ splenocytes. **e-f)** Flow cytometric analysis of cytokine production by CD8^+^ (e) and CD4^+^ T cells (f) in perfused lung tissue. Representation same as b-c. **g-i)** Serum anti-OVA pan-IgG (g), IgG1 (h), and IgG2c (j) titers were determined against OVA protein. **j)** Ratio of IgG2c:IgG1 antibody response from h) and i). Mock vaccines contained antigen only. Values depicted are means±SD. *p < 0.05, **p < 0.01, ***p < 0.001, ****p < 0.0001 by one-way ANOVA followed by Tukey’s post-hoc analysis. n.d., not detected.

### In NHP, AMP-dT_50_ induces cross-protective cellular and humoral immune responses to multiple variants of concern

Encouraged by the immune responses elicited by AMP-DNA in mice, AMP-dT_50_ was assessed in rhesus macaque NHPs for activity predictive of potential responses in humans using a relevant SARS-CoV-2 model. Three adult, female macaques were immunized twice with WH-01 RBD protein admixed with AMP-dT_50_ at week 0 (baseline, BL) and week 4; PBMC, serum, and LN fine-needle aspirates (FNA) were collected for analysis of cellular and humoral immunity as illustrated in Fig. 4a. At week 6, two weeks after the booster dose, significant increases in the frequency of IFNγ SFC were observed among peripheral blood T cells upon restimulation with WH-01 RBD overlapping peptides (OLP) compared to pre-vaccination baseline values in all three animals (Fig. 4b). Moreover, CD4^+^ T cells exhibited significantly increased production of IFNγ, TNFα and IL2 cytokines, and responses in CD8^+^ T cells were elevated compared to baseline (Fig. 4c,d). Similar cellular responses were observed when PBMCs from WH-01 immunized animals were restimulated with OLPs to SARS-CoV-2 variants of concern (VOC), Beta and Delta RBD (Fig. S7a-c), suggesting that AMP-dT_50_ is able to generate robust cellular responses to conserved epitopes potentially important for protection across different strains of SARS-CoV-2. To investigate the humoral response, serum samples were collected in 2-week intervals starting at pre-treatment baseline (Fig. 4e). WH-01 RBD-specific IgG levels were assessed by ELISA and converted into WHO binding antibody units (BAU)/ml. All three animals achieved seroconversion with significantly increased anti-RBD-specific IgG concentrations after only one immunization and reached peak levels two weeks after the second dose. Elevated antibody titers were maintained throughout the assessed study period. When examining antibody responses to Beta, Delta and Omicron RBD variants, the same trends were observed (Fig. S7d,e), substantiating the notion that AMP-dT_50_ adjuvants may generate cross-protective immunity to multiple VOC. To ascertain whether these antibody responses were able to provide neutralizing immunity, which protects against SARS-CoV-2 before infection occurs, pseudovirus inhibitory activity was assessed (Fig. 4f; S7f,g). The antibody response in all animals provided neutralizing protection against the tested VOC at significantly greater ID_50_ values than convalescent human plasma (CHP) from patients that had recently recovered from SARS-CoV-2 infection, suggesting that AMP-dT_50_-induced immunity may exceed response levels generated by natural infection in humans. This humoral response directly correlated with robust germinal center (GC) formation in these animals as determined by flow cytometric analysis of GC B cell (CD20^+^ BcL6^+^ Ki-67^+^) specificity to the RBD antigen in LN FNA (Fig. 4g), a prerequisite to affinity maturation and generation of highly specific antibodies. The immunogenicity of this AMP-dT_50_ adjuvanted SARS-CoV-2 vaccine was accompanied by promising safety assessments; no adverse effects were observed in clinical evaluations (body temperature, weight, injection site observations; Fig. S8a-b), serum cytokine levels (Fig. S8c), and complete blood cell count (Fig. S8d-h), with only a transient increases in select serum cytokines (IFNγ, IL1RA, IL-18) upon administration of AMP-dT_50_ that subsided after 1-2 days. Together, the strong cellular and humoral responses and lack of adverse effects observed in NHP, combined with clinical data using a similar AMP-conjugated CpG adjuvant^27^, indicates promise for AMP-DNA adjuvant evaluation in future clinical studies.

**Figure 4:**
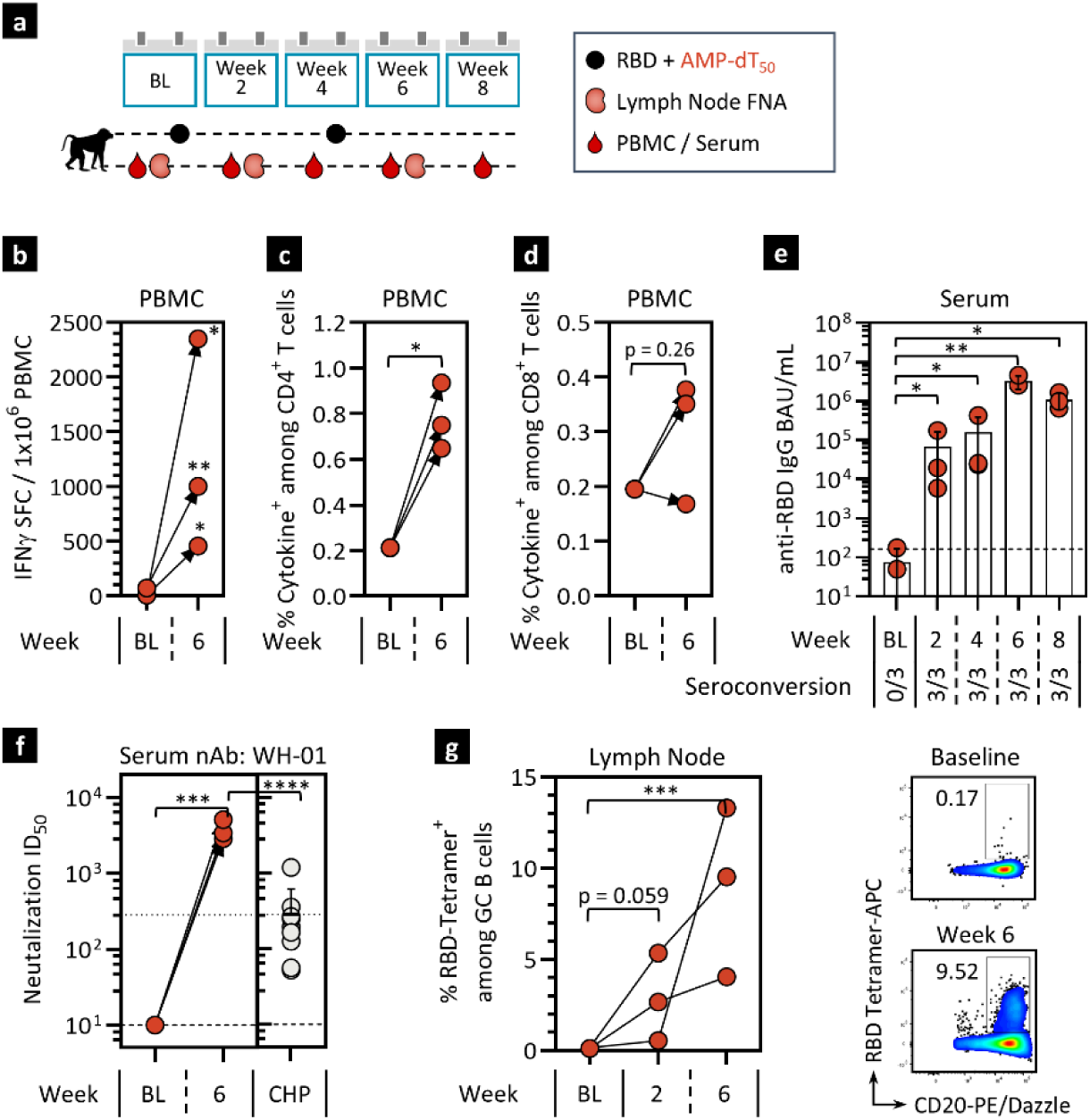
AMP-dT induces potent cellular and humoral immunity in non-human primates. Cellular and humoral responses to WH-01 RBD and AMP-DNA adjuvant were assessed in Rhesus macaques. **a)** 3 animals were immunized at week 0 (Baseline, BL) and 4 with 140 µg WH-01 RBD protein admixed with 5 mg of AMP-dT_50_. **b-d)** PBMCs were collected at BL and week 6 and stimulated with WH-01 RBD OLPs overnight. b) IFNγ ELISpot analysis. Shown are IFNγ SFCs per 1 x 10^6^ PBMCs. c, d) Flow cytometric analysis of cytokine production by CD4^+^ and CD8^+^ T cells. Shown are frequencies of combined IFNγ, TNFα and IL2 cytokine production. **e)** For antibody response assessment serum was collected at BL and weeks 2, 4, 6, and 8. RBD-specific serum IgG binding antibody units (BAU) were assessed for WH-01 at each time point. **f)** For neutralizing antibody (nAb) response assessment through pseudovirus inhibition serum was collected at BL and week 6. Shown are ID_50_ values for WH-01 in comparison to convalescent human plasma (CHP). The dashed lines in e-f) represent the lower limit of detection discriminating between samples positive or negative for seroconversion. The dotted line in f) represents the mean value observed for the human plasma comparators. **g)** For GC B cell assessment LN fine needle aspirates (FNA) were collected at BL and weeks 2 and 6. Shown are frequencies of RBD-specific GC B cells (CD20^+^ BcL6^+^ Ki-67^+^) in individual animals over time with corresponding representative flow cytometry scatter plots of RBD-specific B cells. Values depicted are mean±SD. *\* p < 0.05; ** p < 0.01; *** p < 0.001; **** p < 0.0001* by paired t test.

### AMP-DNA promotes a highly immunostimulatory milieu in lymph nodes of mice and NHP

The improved immunogenicity of AMP adjuvants has been correlated with their enhanced delivery to draining LNs^22,23^. To elucidate the mechanisms within LNs that facilitate AMP-DNA-induced immunity, a comprehensive panel of >500 immune-relevant gene transcripts was evaluated in mice and NHP using NanoString. Mice were immunized once with WH-01 RBD antigen and dT_50_ DNA; subsequently, inguinal LN were collected for transcriptome assessment at the indicated times (Fig. 5a). After 6 hours, mice treated with AMP-dT_50_ displayed an acute upregulation of dozens of transcripts essential to mounting an effective innate immune response (Fig. S9a-c). Upregulated genes included chemokines associated with APC recruitment to the LNs, APC lineage and activation markers, PRRs, and transcripts that facilitate antigen processing and presentation, consistent with professional APC influx and subsequent robust innate immunity. In contrast, mice treated with SOL dT_50_ displayed a much more restricted transcriptional signature limited to a few chemokines. 24 hours after AMP-dT_50_ immunization, the initial innate immune response developed into a fully matured proinflammatory milieu, characterized by multiple axes of immune pathway activation critical to adaptive immunity, while SOL dT_50_ failed to do so (Fig. 5b-d; S9d). Of particular interest were gene profiles indicating enhanced immune functions of activated APCs to recognize danger signals (e.g. *Tbk1, Ddx58* [RIG-I], *Ifih1* [MDA5], *Tlr3, Tlr9, MyD88*), process and present antigens (e.g. *H2-k1, Tap1, B2m, Psmb9/10, Ctss, Ctsc*), co-stimulate T cells (e.g. *Cd80, Cd86, Cd40*), signal via cytokines (e.g. *Tnfa, Il18, Stat2/3, Jak2*) and induce numerous anti-viral interferon-stimulated genes (ISG) such as from the *Irf-*, *Ifi-* and *Ddx*-family of proteins, among many others. At 72 hours post administration of AMP-dT_50_ this inflammatory response had subsided, with PRRs as well as cytokine and chemokine levels returning to baseline (Fig. S9e-g). Nonetheless, a large contingent of cells expressing CD11b (*Itgam*) transcripts remained in the LN. At these early timepoints, no elevation of transcripts associated with increased overall T cell populations were observed relative to untreated groups. Of note, the transcriptome of SOL dT_50_ treated LNs started to develop B cell associated signatures (*Cd19/22/79a/81, Btk, Pax5*) at the 72-hour time point (Fig. S9e), corresponding with the predominant humoral response observed in these animals (Fig. 1h).

**Figure 5:**
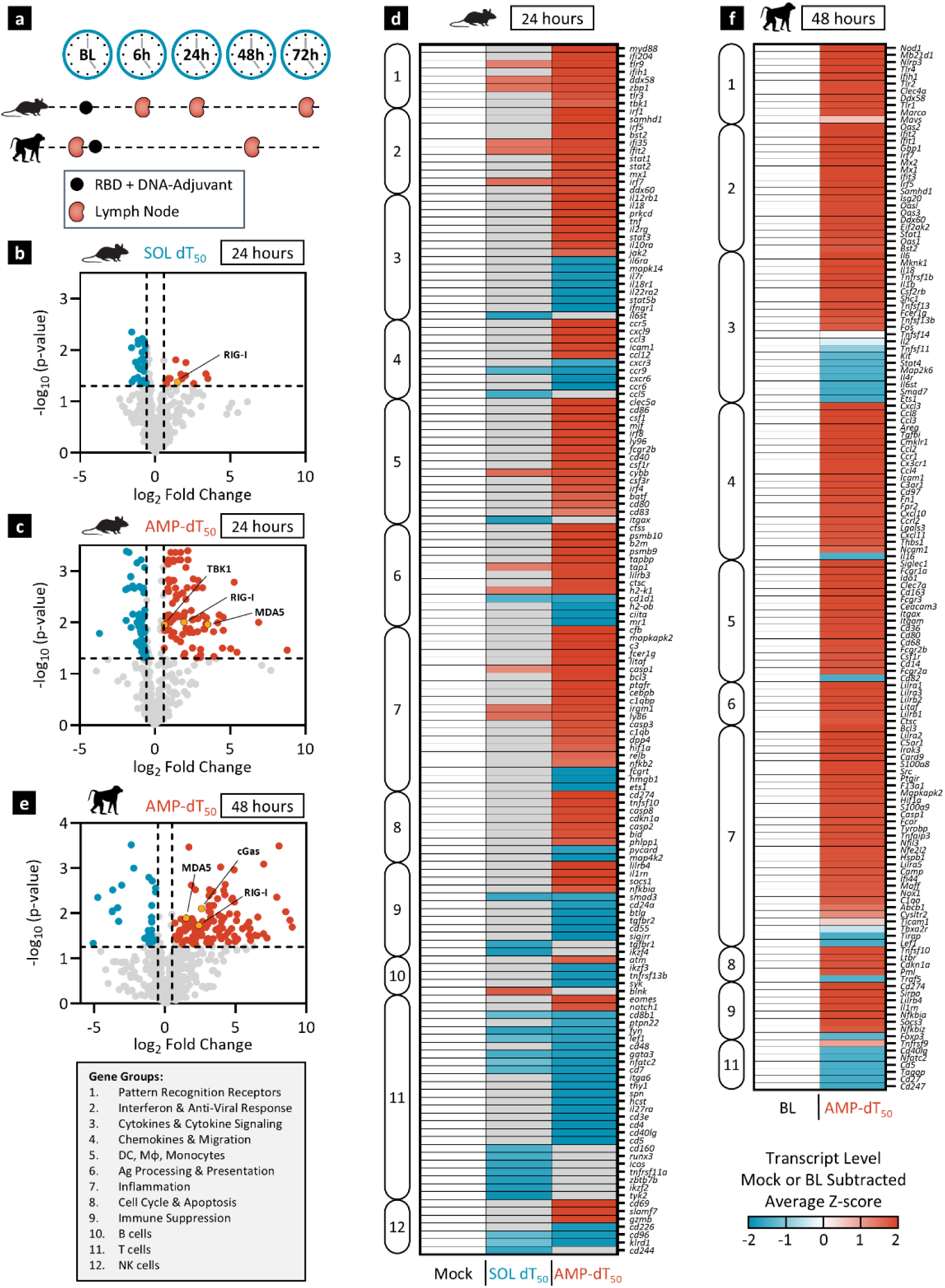
AMP-DNA promotes comprehensive inflammatory transcriptional reprogramming in draining lymph nodes of mice and NHP. **a)** Schema showing animal dosing and experimental schedule. C57BL/6J mice (n = 2) were immunized once with 5 μg WH-01 RBD protein admixed with 5 nmol SOL or AMP-dT_50_ adjuvants. Whole LN lysates were assayed by NanoString nCounter Mouse Immunology Panel at the indicated time points post primer dose. Rhesus macaques (n = 2) were immunized once with 140 µg WH-01 RBD protein admixed with 5 mg of AMP-dT_50_ adjuvant. LN FNAs were collected prior to week 10 boost (baseline) and 48 hours after booster immunization and analyzed by NanoString nCounter® NHP Immunology Panel. **b-c)** Volcano plot representation of log-transformed transcript values from SOL dT_50_ (b) and AMP-dT_50_ (c) immunized mice at 24 hours (red values are significantly upregulated and blue values are significantly downregulated gene transcripts that changed by a minimum of 1.5-fold). Dashed horizontal line represents significance threshold (p = 0.05); vertical dashed lines represent fold-change limits of ±1.5. **d)** Heatmap representation of transcript expression 24 hours after immunization. Shown are mock-subtracted, average Z-scores of transcripts that are significantly changed in at least one treatment group and that are either ≥1.5-fold upregulated (red) or downregulated (blue) compared to Mock treatment. Mock vaccines contained PBS vehicle only. Insignificant values (p ≥ 0.05) are shown in gray. **e)** Volcano plot representation of log-transformed transcript values from AMP-dT_50_ immunized NHP at 48 hours after immunization (significance and fold-change thresholds as in b,c). **f)** Heatmap representation of LN FNA mRNA analyzed 48 hours after immunization. Shown are baseline-subtracted, average Z-scores of transcripts that are significantly changed and are either ≥1.5-fold upregulated (red) or downregulated (blue) compared to baseline. Genes were clustered into 12 groups (box insert) using Gene Ontology and UniProt databases.

Similar results were observed in NHPs upon immunization with WH-01 RBD admixed with AMP-dT_50_. Transcriptome analysis of collected LNs showed the upregulation of multiple chemokines and cell adhesion molecules 24 hours after AMP-dT administration compared to pre-vaccination baseline values, suggesting an influx of innate immune cells (Fig. S10a,b). At 48 hours, this was followed by further augmentation of a wide range of proinflammatory pathways (Fig. 5e,f; S10c,d). The increased transcriptional signature of various APCs in the LN including dendritic cells (DC) and macrophages (*Fcgr3* [CD16], *Fcgr2a/b* [CD32], *Fcgr1a* [CD64], *Itgam* [CD11b], *Itgax* [CD11c]) correlated with a highly anti-viral environment (*Oas1/2/3, Ifit1/2/3, Irf5/7*) sensitive to nucleotide detection (*Ddx58* [RIG-I], *Ddx60*, *Ifih1* [MDA5], *Mb21d1* [cGas], *OasL*). These transcriptional features were consistent with the development of an inflammatory niche supportive of the adaptive immune responses observed with AMP-DNA immunizations in NHP (Fig. 4). Across mice and NHPs, AMP-dT_50_ activated numerous shared immune pathways required to convert the naïve LN into a proinflammatory hotspot (Fig. S10e). Correlation of these transcriptional patterns to immunogenicity further highlights the importance of engaging the innate immune system in this specialized organ, which is an optimal environment for robust immune activation.

### AMP-DNA potently induces inflammatory cytokine responses dependent on IFN-I signaling

Given the notable induction of immunostimulatory transcriptional programs upon vaccination with AMP-DNA, we investigated if these effectively translated into changes in the proteomic profile within the LNs. Mice were immunized with one dose of WH-01 RBD and AMP-dT_50_ (Fig. 6a; S11) or AMP-dA:dT_50_ (Fig. S12) or their respective soluble counterparts and draining LNs were collected 6 or 24 hours later for multiplexed proteomic analysis. AMP-dT_50_-treated animals exhibited a significant boost in the production of nearly all tested cytokines over mock- or SOL-dT_50_-treated mice at the 24-hour time point, consistent with mRNA transcript levels at that time. These cytokines included 1) growth factors (G-CSF, GM-CSF, M-CSF), 2) proinflammatory cytokines (IFNγ, TNFα, IL6), 3) chemokines (IP-10, IL8, MCP1, MIP1α/β, MIG), 4) inflammasome-associated cytokines (IL1β, IL18), and 5) IFN-I, consistent with the ISG-signature observed above. Similarly, mice immunized with AMP-dA:dT_50_ had high levels of cytokine production at 24 hours but appeared to initiate production earlier than the single stranded form. In contrast, cytokine levels in SOL dA:dT_50_-treated LNs were significantly lower than in AMP groups, and despite exhibiting detectable cytokine upregulation over baseline, did not reach sufficient levels to induce effective downstream immune induction (Fig. 1).

**Figure 6:**
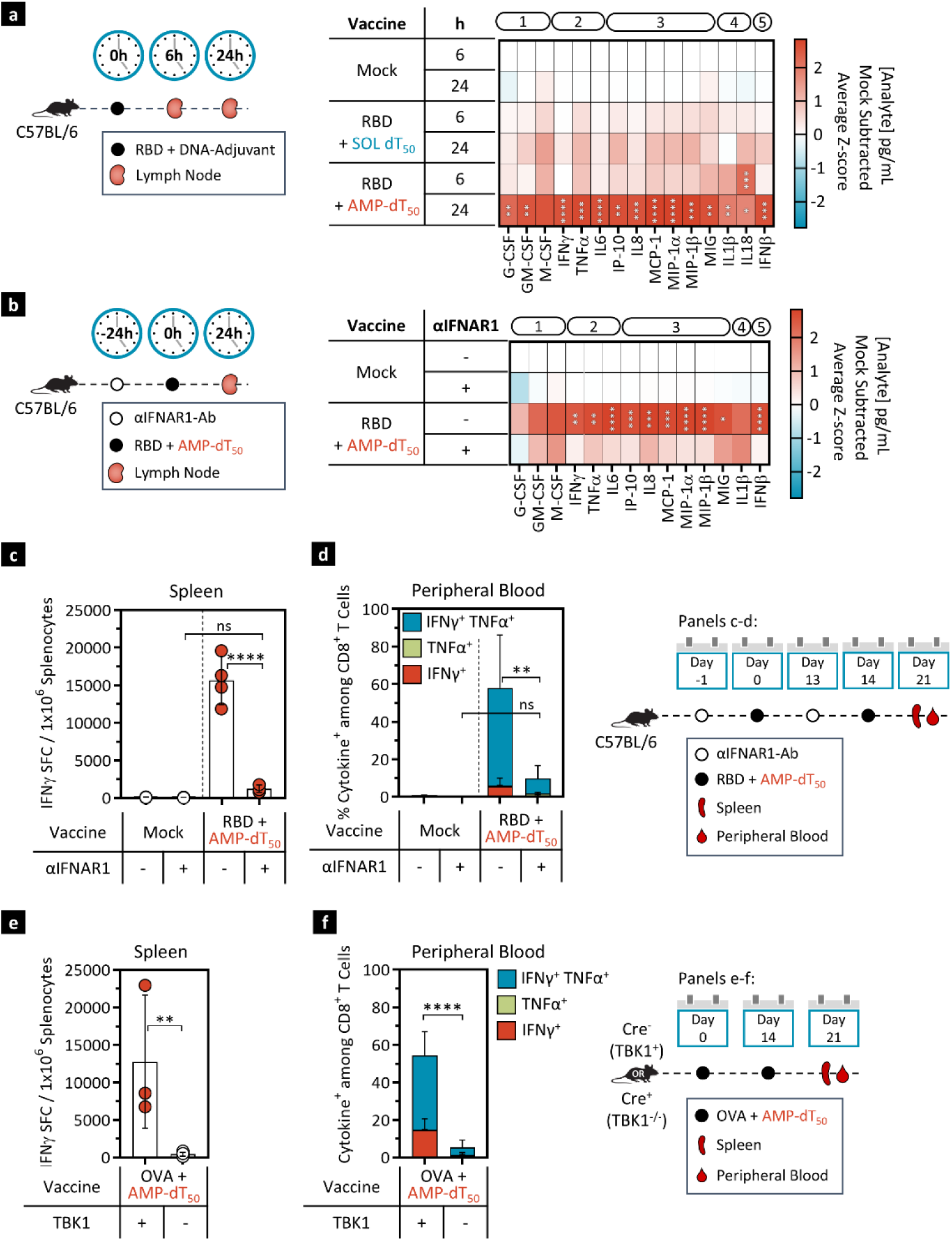
AMP-dT_50_ potently induces an inflammatory proteomic milieu in draining lymph nodes dependent on IFN-I and TBK1. **a)** C57BL/6J mice (n = 5) were immunized once with 5 μg WH-01 RBD protein and 5 nmol SOL or AMP-conjugated dT_50_. Lymph nodes were collected and processed for protein extraction and analysis by Luminex at 6 and 24 hours post immunization. Analyte concentrations were expressed in a heat map showing mock-subtracted Z-scores of analyte concentrations (pg/ml). Analytes are annotated by functional category: (1) growth factors, (2) inflammatory cytokines, (3) chemokines, (4) inflammasome, and (5) IFN-I. Statistical analysis compares AMP-DNA treated groups with time point-matched SOL DNA groups. **b)** C57BL/6J mice (n = 5) were dosed intraperitoneally with IFNAR1 blocking antibody or isotype control 24 hours prior to immunization as in a). Protein from lymph nodes was extracted 24 hours post immunization. Heatmap representation and analyte annotation as in a). Statistical analysis compares IFNAR1-Ab treated groups with isotype-Ab treated groups. **c-d)** C57BL/6J mice (n = 5) were immunized twice with 5 μg WH-01 RBD protein and 5 nmol AMP-dT_50_. Mice received IFNAR1 blocking antibody or isotype control intraperitoneally one day prior to each dose. Cells were assayed 7 days post booster dose. c) Splenocytes were restimulated with RBD OLPs overnight and assayed for IFNγ production by ELISpot. Shown is the frequency of IFNγ SFCs per 10^6^ splenocytes. d) Flow cytometric analysis of cytokine production by CD8^+^ T cells in peripheral blood upon restimulation with RBD OLPs. Mock immunized animals received PBS vehicle only. **e-f)** TBK1^fl/fl^ x Vav-iCre^+/-^ mice (TBK1^-/-^, n = 8) and TBK1^fl/fl^ x Vav-iCre^-/-^ mice (TBK1^+^, n = 5) were immunized twice with 5 μg OVA protein and 5 nmol AMP-dT_50_. Cells were assayed 7 days post booster dose. e) Splenocytes were restimulated with SIINFEKL peptide overnight and assayed for IFNγ production by ELISpot. Shown is the frequency of IFNγ SFCs per 10^6^ splenocytes. f) Flow cytometric analysis of cytokine production by CD8^+^ T cells in peripheral blood upon restimulation with SIINFEKL peptide overnight. Values depicted are mean±SD. *p < 0.05; **p < 0.01; ***p < 0.001; ****p < 0.0001 by one-way ANOVA followed by Tukey’s post-hoc analysis.

Of particular note is the strong AMP-DNA-mediated induction of IFN-I, an important mediator of robust immune activation following initial pathogen detection^34,35^. To investigate its impact on AMP-DNA-induced immune activation, IFNAR1, essential for IFN-I signaling, was blocked in mice using an anti-IFNAR1 antibody 24 hours prior to immunization with WH-01 RBD and AMP-dT_50_ (Fig. 6b). This resulted in nearly complete abrogation of the consequent immune response. Assessment of cytokine levels in draining LNs revealed a significant reduction in almost all analytes measured compared to mice treated with an isotype control antibody (Fig. 6b; S13a-d), leading to a significantly attenuated T cell response in spleen, peripheral blood and perfused lung (Fig. 6c-d; S13e-g). Only the humoral response was relatively unaffected by IFNAR1 blockade as determined by antibody titers to WH-01 RBD (Fig. S13h). Similar results were observed in mice immunized with AMP-dA:dT_50_ (Fig. S14). Overall, these data demonstrate that AMP-DNA adjuvants induce a LN environment rich in numerous pro-inflammatory cytokines, of which IFN-I plays a critical role in conducting the AMP-DNA signal leading to adaptive immunity.

### AMP-DNA adjuvants require TBK1 signaling

Immune responses are strongly induced upon nucleic acid detection by cytosolic PRRs^35^. These sensors initiate pathways that either form inflammasomes, which control the proteolytic maturation of IL1 and IL18, or that converge on TBK1 to induce transcription of IFN-I. The two major TBK1-activating pathways reside in the cytosol and include DNA-sensing and RNA-sensing receptors which signal through cGAS-STING, and RIG-I/MDA-5/MAVS, respectively^36,37^. To determine if AMP-DNA-induced immune activation relies on TBK1 signaling, TBK1-floxed mice expressing Cre-recombinase under the hematopoietic cell-specific *Vav1*-promoter were immunized twice with OVA and AMP-dT_50_ (Fig. 6e,f). Mice with TBK1-deletions in the hematopoietic compartment (TBK1^fl/fl^ x Vav-iCre) displayed a significantly diminished ability to generate splenic IFNγ SFCs and showed significantly reduced frequencies of circulating cytokine^+^ CD8^+^ T cells, indicating that AMP-DNA requires TBK1 signaling to induce cellular immune responses. Overall, these data suggest that AMP-DNA adjuvants must gain access to the cytosol of immune cells to engage cytosolic PRRs that signal through TBK1 and induce potent IFN-I responses in APCs that ultimately lead to effective cellular immunity complementing a robust humoral response.

## DISCUSSION

Highly purified, recombinant subunit antigens yield remarkably safe and precise, albeit often poorly immunogenic vaccines that require the presence of potent adjuvants to enhance their immunogenicity. Notable un-adjuvanted exceptions include virus-like particle vaccines made from proteins that self-assemble into multimeric and highly immunogenic viral structures^38^; conjugate vaccines that link polysaccharide antigens to carrier proteins such as tetanus or diphtheria toxoid that support T cell helper functions^39,40^, and, rarely, antigens that are sufficiently immunogenic to generate appropriate immune responses unaided^41,42^. However, responses to most of these vaccine classes are predominantly antibody-mediated and immunity to the latter is improved by the addition of adjuvant^43,44^. Therefore, the next generation of adjuvant systems will need to combine several important capabilities in order to be most effective: 1) In order to increase bioavailability and durable immune activation while minimizing reactogenicity, vaccines will benefit from a targeted approach that delivers adjuvants directly to LNs where critical immune cells reside; 2) To induce potent immunostimulation, especially activation of IFN-I crucial for antiviral and antitumor immunity^45,46^, adjuvants will need to access endosomal TLRs or cytosolic PRRs, which are known to strongly induce these activation pathways^4^; 3) Lastly, a balanced cellular and humoral immune response is critical to achieve comprehensive immunity. To generate robust CD8^+^ cytotoxic T lymphocyte (CTL) responses, new adjuvants must promote cross-presentation within APCs^47^. In this study, we have engineered a class of DNA-adjuvants, most prominently represented by AMP-dT_50_, that exhibit all three of these traits, exemplifying the potential gains in activity and tolerability that AMP-modification can provide.

Consistent with prior studies, AMP-modification of candidate DNA sequences resulted in efficient and targeted delivery to draining LNs corresponding with improved immunogenicity and favorable safety. In contrast, most immunostimulatory PRR/TLR agonists, such as small molecules or short nucleic acid sequences, fail to efficiently accumulate in immunorelevant tissues because of generally disadvantageous physicochemical characteristics (e.g.: size <20 kDa)^13^. This results in poor immunogenicity despite their inherent potential to activate the innate immune system. Consequently, delivery platforms are required to increase bioavailability and in turn, potency. AMP-conjugation of immunostimulatory adjuvants provides a simple and versatile strategy to guide vaccine components via tissue albumin-mediated shuttling from the injection site directly to draining LNs where they activate APCs^22,23^. Moreover, the association of AMP-constructs with albumin is known to protect payloads from degradation and prolong bioavailability^23,48^. Here, we have demonstrated that AMP-conjugation significantly enhances immunostimulatory properties of candidate DNA-adjuvants, such as dT_50_, dA:dT_50_, HSV_60_, and ISD upon prime-boost administration. In mice, production of cytokines from splenic, lung-resident, and circulating antigen-specific T cells was increased multiple dozen- to hundred-fold when compared to SOL DNA adjuvants or clinical adjuvant comparators, which achieve limited LN accumulation. In parallel, antibody responses to AMP-DNA equaled that of classical depot-forming adjuvants which are known to primarily elicit humoral responses. However, AMP-DNA adjuvants produced stronger T_H_1-associated IgG_2c_ titers that likely play a supportive role in facilitating the significant cellular response present in these animals. Further, NHPs treated with AMP-dT_50_ and RBD antigen exhibited neutralizing antibody titers that exceeded those of convalescing patients who had recently recovered from SARS-CoV-2 infection. The notable immunogenicity-enhancing properties of these AMP-adjuvants align with previous reports using AMP-CpG-7909 in vaccines targeting SARS-CoV-2^25,26^ and EBV^49^. Furthermore, a recent clinical trial with AMP-CpG-7909 in KRAS-mutated pancreatic and colorectal cancer patients^27^ showed that targeted delivery of vaccine components can promote favorable immunogenicity while avoiding systemic toxicity. Treatment-emergent adverse events were limited to grade 1-2; no dose-limiting toxicity or cytokine-release syndrome was observed. Concurrently, 84% of patients overall and 100% of those treated at the recommended phase 2 dose, showed mKRAS-specific T cell responses *ex vivo* which strongly associated with tumor biomarker responses and decreased risk of progression and death. Overall, these data demonstrate that targeted delivery not only increases the potency of vaccines by concentrating their components in immunorelevant tissues but can also prevent harmful side effects.

Secondly, for adjuvants to be effective immunostimulators, they must activate APCs properly by gaining access to and interacting with the appropriate PRRs within the cell. Responses against cancers^50^ and intracellular pathogens such as viruses^51^, tuberculosis^52^ and malaria^8,53^ rely partly on cellular immune responses that are profoundly benefited by localized IFN-I production in the LN. Several cytosolic PRRs are strong inducers of such IFN signaling pathways^35^ as they evolved to detect nucleic acids from intracellular pathogens or self-DNA from compromised or stressed cells^54^. APCs contain numerous cytosolic nucleic acid sensors, which detect either DNA or RNA and signal through the STING or MAVS pathways, respectively, culminating in the activation of both TBK1 and NFκB to ultimately initiate the secretion of IFN-I and other inflammatory cytokines that drive the cellular adaptive immune response^45,46^. This makes these APC-associated, nucleic acid-sensing PRRs desirable targets for new adjuvants. In this study we have engineered phosphorothioate-stabilized DNA and explored various design characteristics to optimally stimulate the local immune system within the LNs. This was demonstrated by the significant induction of IFN-I and inflammatory transcripts in the LNs, including multiple cytosolic PRRs and TBK1, antigen processing and presentation machinery, APC activation markers and co-stimulatory receptors, as well as the subsequent increased secretion of various cytokines and chemokines. Most notably, IFN-I played a pivotal role in initiating adaptive immunity, as IFNAR1-blockade abrogated all cellular responses, in agreement with previous studies^45^. Crucially, however, to reach these cytosolic receptors, endocytosed adjuvants must first escape the endocytic pathway and enter the cytosol to avoid entrapment and eventual degradation within the lysosome; a rate-limiting step that determines the efficacy of many therapeutics^55^. The diacyl lipid constituent of the AMP-modification could enable such mechanisms to allow AMP-DNA adjuvants to engage cytosolic TBK1-mediated pathways. AMP-constructs have demonstrated the ability to insert into lipid bilayers or form micellar structures^22^, either of which could potentially facilitate the disruption of endosomal membranes and/or enhance initial APC uptake preceding endosomal escape^56^. The exact mechanisms of AMP-facilitated access to the cytosol require further investigation; however, SOL dT_50_ was unable to initiate notable immune responses, despite sufficiently accumulating in the LNs during fluorescent *in vivo* imaging studies, suggesting that AMP-modification is critical for successful immune induction after the adjuvant accumulates in the LNs. Overall, the data suggest that AMP-DNA adjuvants are strong inducers of critical IFN-I signals needed to promote cellular immunity and that AMP-modification not only facilitates delivery to the LNs but may enhance conjugated vaccine payload entry into the cytosol of immune cells to enable efficient stimulation of PRRs.

Finally, improved vaccine potency depends in part on cross-presentation of antigens to generate robust populations of cytotoxic effector and memory T cells. Cellular immunity is an essential component of protection against infected or transformed cells and operates in synergy with a parallel humoral response. However, most vaccines in use today, that do not contain live-attenuated or inactivated (self-adjuvanting) vectors, primarily elicit antibody responses^5,57,58^. In isolation, these are often insufficient to provide adequate protection against diseases caused by many intracellular pathogens or cancer, which instead require combined humoral and cellular immunity^59^. For example, neutralizing antibodies were not sufficient to protect patients against fatal SARS-CoV-2 infection, suggesting that cellular responses are required for comprehensive protection against this virus^60^. Moreover, antibody responses in the presence of inadequate innate immune activation and thus lack of T helper cell participation are generally short-lived; most responses to SARS-CoV-2 vaccine begin to wane significantly within 3-6 months after vaccination^61–63^. Thus, to achieve appropriate APC activation that stimulates potent humoral responses, while simultaneously promoting robust and durable cellular immunity, new adjuvants will ideally induce strong IFN-I signaling, which enhances their intrinsic cross-presentation and co-stimulatory capabilities^64,65^, resulting in strong CD8^+^ T cell priming and memory generation^18,66^. AMP-DNA adjuvants demonstrated effective activation of antigen-specific CD8^+^ T cell populations in secondary lymphoid and peripheral tissues as well as in circulation. This was accompanied by significantly elevated production of IFN-I, and the enhanced transcription of genes important for cross-presentation. Of note, transporter associated with antigen processing 1 (*Tap1*), TAP-binding protein (*Tapbp*) and proteasome subunits (*Psmb9/10*), all crucial in the cytosolic pathway of cross-presentation^64^, as well as cathepsins (*Ctss, Ctsc*) involved in the vacuolar pathway of cross-presentation^67,68^, were transcriptionally upregulated in LNs upon vaccination with AMP-DNA adjuvants. Together these findings indicate an ability of AMP-DNA adjuvants to potently induce cellular immune responses facilitated by significant production of IFN-I and the promotion of cross-presentation of antigens via MHCI molecules.

We are aware of some limitations in the current study. While immunogenicity data obtained from NHP (n=3) corroborate the notable responses obtained in mice and suggest the potential of AMP-DNA as an additional example of efficacious and safe AMP-adjuvant systems for use in humans, larger studies will be useful to support more decisive conclusions about its potential clinical utility. While cytosolic TBK1 is involved in signal transduction of AMP-DNA adjuvants, these initial findings will need to be substantiated by more detailed analysis of cellular uptake and signaling pathways, including assessment of STING, MAVS, TLRs, and the inflammasome, to precisely define how AMP-DNA adjuvants are sensed and how they shape these downstream mechanisms. Further investigation is also needed to determine the mechanisms of cytosolic entry and participation of critical immune cell lineages. Moreover, immunization studies, including the extended duration cohort which was analyzed at 3 months post vaccination, were analyzed at relatively short timepoints. Thus, conclusions about the durability of the AMP-DNA-induced immune response will benefit from studies monitoring cellular and humoral immunity over more extensive time periods. Lastly, this study is currently limited to immunogenicity data upon vaccination with AMP-DNA adjuvants and should be complemented by further pathogen or tumor challenge studies that can determine the adjuvant effect on vaccine efficacy.

In summary, AMP-DNA represents a novel class of potent and safe immunostimulatory adjuvants that generate significant humoral and cellular immunity to co-administered protein subunit antigens. Strong immunogenicity is associated with adjuvant design parameters facilitating direct targeting of the innate immune system within LNs, stimulation of cytosolic PRR pathways, and induction of potent IFN-I responses. These and previous findings suggest that AMP-conjugated adjuvants such as AMP-DNA and AMP-CpG-7909, are promising candidate vaccine adjuvants for clinical use in infectious disease and cancer.

## METHODS AND MATERIALS

### Animals and tissue processing

All animal studies were carried out under a New Iberia Research Center- or Charles River Accelerator and Development Lab-approved Institutional Animal Care and Use Committee (IACUC) protocol following federal, state, and local guidelines for the care and use of animals. For mouse studies, female 6- to 8-week-old C57BL/6J mice were purchased from the Jackson Laboratory (Bar Harbor, ME). B6-TBK1^fl/fl^ mice were generated by Millennium Pharmaceuticals (Takeda) and crossed with Vav-iCre mice to generate offspring with TBK1 deletions specifically in the hematopoietic compartment. The indicated antigen-adjuvant combinations were administered into mice subcutaneously at the base of the tail (bilaterally, 50 µl each) at weeks 0 and 2. Spleen and perfused lung samples were collected 7 days post booster dose. Spleens were processed into single-cell suspensions and red blood cells were lysed in ACK lysis buffer (Quality Biological Inc., cat# 118156101). Lungs were harvested following perfusion with 10 mL of PBS into the right ventricle of the heart. Perfused lung tissue was physically dissociated and digested with RMPI 1640 media containing collagenase D (1 mg/mL) and deoxyribonuclease I (25 U/mL). Peripheral blood samples were collected retro-orbitally at the indicated time points and processed for PBMCs and serum. LN samples were collected at 2, 6, 24, and 72 hours post primary dose for transcriptome analysis, and 6 and 24 hours post primary dose for proteomic analysis.

For NHP studies, animals were housed at New Iberia Research Center (New Iberia, LA). 3 outbred, Indian-origin, 4-5 year old female rhesus macaques (*Macaca mulatta*) received immunizations subcutaneously into the upper thigh at week 0 and 4 for evaluation of T cell and antibody responses. Blood samples were obtained at baseline and throughout the study in two-week intervals and were then processed for PBMCs and serum. LNs were sampled via FNA at baseline, week 2 and 6, and analyzed for GC B cell specificity. For analysis of the inguinal LN transcriptome, FNAs were collected 24 and 48 hours after a third dose at week 10.

### Vaccine components

For mouse studies, vaccines consisted of 5 µg SARS-CoV-2 Spike RBD WH-01 (RBD) protein (GenScript, cat# Z03483) or 5 µg ovalbumin (OVA) protein (InvivoGen, cat# vac-pova), combined with 5 nmol DNA adjuvant. The adjuvants included AMP-dA:dT, AMP-dT, and any of their length, sequence, or linkage variants as well as their soluble counterparts (SOL dA:dT and SOL dT) as indicated. AMP-CpG-7909 was used at 1 nmol per injection. All were synthesized by AxoLabs GmbH (Germany). Comparator adjuvants included Alhydrogel adjuvant 2% (Alum, 100 µg, InvivoGen, cat# vac-alu), AS03-like squalene-based adjuvant (AddaS03, 1:2 dilution, InvivoGen, cat# vac-as03), Monophosphoryl Lipid A (MPLA, 10 µg, InvivoGen, cat# vac-mpla), MF59-like adjuvant (AddaVax, 1:2 dilution, InvivoGen, cat# vac-adx), and Incomplete Freund’s Adjuvant (IFA, 1:2 dilution, InvivoGen, cat# vac-ifa). For fluorescent *in vivo* imaging studies DNA-adjuvants were 3’-conjugated to Cy5 fluorophore (AxoLabs). Mock treatment groups received a matching dose of antigen in the absence of adjuvant or were treated with vehicle alone (phosphate-buffered saline, PBS), as indicated. For NHP studies with Rhesus macaques, vaccines consisted of 140 µg SARS-CoV-2 Spike RBD WH-01 protein admixed with 5 mg of AMP-dT_50_.

### DNA-adjuvant synthesis

Single stranded phosphorothioated 2′-deoxyoligonucleotides were synthesized according to the conventional solid-phase oligonucleotide synthesis technology using standard phosphoramidite-based oligomerization chemistry. Oligonucleotides were assembled on solid support employing a Mermade 12 synthesizer (LGC Bioautomation) controlled by the Poseidon software package (version 2.7.0.0). In order to introduce phosphorothioate linkages a 100 mM solution of 3-Amino-1,2,4-dithiazole-5-thione dissolved in ACN-pyridine (2:3 v/v) was employed as sulfurizing agent. Diacyl-lipid phosphoramidite coupling was performed using ETT (500 mM) formulated in isobutyronitrile-DCM (1:1 *v/v*). Oligonucleotides were cleaved from the solid support with concomitant removal of all protecting groups using AMA (1:1 (v/v) mixture of concentrated aqueous ammonia and 40% aqueous methylamine) with shaking at 25 °C. Samples were then dried under reduced pressure (SpeedVac concentrator (ThermoFisher)) and were finally reconstituted in 100 mM aqueous triethylammonium acetate (TEAAc) to yield crude sample solutions for subsequent purification. Crude preparations were purified by preparative RP-HPLC using a XBridge BEH C18 (19×50 mm, OBD, 5 µm particle) column available from WATERS on an ÄKTA Pure 100 system (Cytiva). Eluent A was 100 mM triethylammonium acetate (TEAA) (pH 7) in water and eluent B was a formulation of eluent A in 95% acetonitrile-water. UV traces at 260 and 280 nm were recorded to monitor product elution. Appropriate fractions were pooled and precipitated overnight in the freezer using 3M NaOAc, pH=5.2 in ethanol (1:32 v/v). Pellets were collected by centrifugation and reconstituted in purified water. DMT-On oligos were thereafter detritylated by treatment with 10% acetic acid, followed by repurification on the same RP-HPLC system. The purified complementary DNA single strands were mixed in an equimolar ratio and annealed by heating to 70 °C for 5 min followed by cooling to 25 °C in 2 h. The resultant DNA duplexes were characterized by non-denaturing HPLC to evaluate their purity (duplex purity > 90%, as per integration of the UV signal at 260 nm of the analytical non-denaturing HPLC trace). HPLC purified DNA single strand equipped with a 3’-aminohexyl-linker was conjugated to a sulfo-Cyanin5 NHS ester (Lumiprobe). Samples were quantified by measuring the UV absorption at 260 nm (based on the theoretical extinction coefficient, computed by the nearest neighbor method and the molecular weight of the sodium salt). Samples were analyzed for purity and identity using LC/MS.

### Re-stimulating Peptides

For *ex vivo* re-stimulation of cells from animals immunized with RBD antigen, custom 15mer OLPs with 11 amino acid overlap were generated spanning the SARS-CoV-2 Spike RBD WH-01 protein (amino acids R319-S591, GenScript). WH-01 peptides that contained VOC mutation loci were substituted with the corresponding mutant sequences when applicable. *Ex vivo* re-stimulation of cells from animals that were immunized with OVA antigen was performed with PepTivator® Ovalbumin OLP (Miltenyi, cat# 130-099-771) or SIINFEKL peptide (AnaSpec Inc., cat# AS-60193-1).

### Cytokine production analysis by flow cytometry to assess T cell activation

For vaccine immunogenicity studies in mice, intracellular cytokine staining was performed for TNFα and IFNγ. Peripheral blood cells and lung-resident lymphocytes were collected 7 days after the booster dose and stimulated overnight (16-20 h) with peptides (2 μg per peptide per ml) at 37°C, 5% CO_2_ in the presence of brefeldin A (Invitrogen, cat# 00-4506-15) and monensin (BioLegend, cat# 420701). A Live/Dead fixable cell stain (Aqua, Invitrogen, cat# L34966) was used to evaluate viability of the cells during flow cytometry. Cells were surface-stained with antibodies against anti-mouse CD4 (PE-Cy7, clone: GK1.5, Invitrogen, cat# 25-0041-82, 1:200 dilution), and anti-mouse CD8a (APC, clone: 53-6.7, eBioscience, cat# 17-0081-83, 1:200) and subsequently fixed and permeabilized with BD CytoFix/CytoPerm (BD, cat# 554714). Cells were further stained intracellularly with antibodies against anti-mouse IFNγ (PE, clone: XMG1.2, BD, cat# 554412, 1:200), anti-mouse TNFα (FITC, clone: MP6-XT22, BD, cat# 554418, 1:200), and anti-mouse CD3 (APC-Cy7, clone: 17A2, BD, cat# 560590, 1:200). PMA (50 ng/ml) and ionomycin (1 μM) were used as positive control, and complete medium plus DMSO (at peptide equivalent volume) was used as the negative control. Sample acquisition was performed on FACSCanto II (BD), and data were analyzed with FlowJo V10 software (BD).

For NHP studies, frozen PBMCs were thawed and rested overnight. 10^6^ PBMCs/well were resuspended in R10 media supplemented with anti-human CD49d monoclonal antibody (clone: 9F10, BD, cat# 555502), anti-human CD28 monoclonal antibody (clone: CD28.2, BD, cat# 555726), and Golgi inhibitors monensin (Fisher Scientific, cat# NC0176671) and brefeldin A (Fisher Scientific, cat# 50-112-9757) and incubated at 37°C for 8 hours, then maintained at 4°C overnight. The next day, cells were surface-stained with antibodies against anti-human CD4 (PE-Cy5.5, clone: S3.5, Invitrogen, cat# MHCD0418, 1:100), anti-human CD8 (AF647, clone: RPA-T8, BioLegend, cat# 301022, 1:400), and Live/Dead dye (Aqua, Invitrogen, cat# L34966), and subsequently fixed with BD CytoFix/CytoPerm (BD, cat# 554714). Cells were further stained intracellularly with antibodies against anti-human CD3 (APC-Cy7, clone: SP34-2, BD, cat# 557757, 1:200), anti-human IFNγ (AF700, clone: B27, BioLegend, cat# 506516, 1:400), anti-human IL-2 (BV421, clone: MQ1-17H12, BioLegend, cat# 500328, 1:20), and anti-human TNFα (BV605, clone: MAb11, BioLegend, cat# 502936, 1:20). Stained cells were fixed in 1.5% formaldehyde and acquired on a BD FACS Symphony. Data were analyzed with BD FlowJo V10 software.

### Mouse antigen-specific tetramer staining to assess antigen-specific T cells

MHC-tetramer staining on mouse peripheral blood samples was performed using an OVA-PE iTAg tetramer specific for sequence SIINFEKL (MBL, cat# TB-5001-1) or RBD-PE tetramer specific for sequence VNFNFNGL (NIH Tetramer Core Facility), and antibodies against anti-mouse CD8a (APC, clone: 53-6.7, eBioscience, cat# 17-0081-83, 1:200), anti-mouse CD3 (APC-Cy7, clone: 17A2, BD, cat# 560590, 1:200). A Live/Dead fixable cell stain (Aqua, Invitrogen, cat# L34966) was used to evaluate viability of the cells during flow cytometry. Sample acquisition was performed on a BD FACSCanto II and data were analyzed with BD FlowJo V10 software.

### NHP antigen-specific GC B cell analysis in lymph node FNAs

Biotinylated RBD protein (Acro Biosystems, cat# SPD-C82E9) was complexed with fluorochrome-conjugated streptavidin APC (BioLegend, cat# 405207). LN cells from FNA were incubated with the RBD-fluorochrome complex and subsequently stained with Live/Dead dye (Aqua, Invitrogen, cat# L34966), anti-human IgM (FITC, clone: G20-127, BD, cat# 555782, 1:5), anti-human IgG (PE-Cy7, clone: G18-145, BD, cat# 561298, 1:20), anti-human CD3 (AF700, clone: SP34-2, BD, cat# 557917, 1:20), anti-human PD-1 (BV650, clone: EH12.1, BD, cat# 564104, 1:20), anti-human CD20 (PE/Dazzle 594, clone: 2H7, BioLegend, cat# 302348, 1:20), anti-human CD4 (APC-Cy7, clone: OKT4, BioLegend, cat# 317418, 1:20) and anti-human CXCR5 (PerCP-eF710, clone: MU5UBEE, ThermoFisher, cat# 46-9185-42, 1:20). Cells were fixed/permeabilized using Transcription Factor Staining Buffer Set (ThermoFisher, cat# 00-5521-00) and further stained with anti-human Bcl-6 (PE, clone: 7D1, BioLegend, cat# 358504, 1:20) and anti-human Ki-67 (BV421, clone: 11F6, BioLegend, cat# 151208, 1:20). Sample acquisition was performed on a BD FACS Symphony and data were analyzed with BD FlowJo V10 software.

### Enzyme Linked ImmunoSpot (ELISpot) Assay for IFNγ to assess T cell activation

For mouse studies, precoated 96-well mouse IFNγ ELISpot plates (MabTech, cat# 3321-4HPW) were blocked with RPMI + 10% FBS for 2 hours at room temperature. 0.1×10^6^ mouse splenocytes were plated into each well and stimulated overnight with RBD- or OVA-derived OLPs (2 µg/ml per peptide). ELISpot assays were performed as instructed by the manufacturer.

For NHP studies, ELISpot assays were performed using Monkey IFNγ ELISpotPLUS kits (MabTech, cat# 3241M-4HPW). Precoated 96-well ELISpot plates were blocked with RPMI + 10% FBS for 2 hours at room temperature. 0.2×10^6^ PBMCs were plated into each well and stimulated overnight with RBD-derived OLPs for WH-01, Beta and Delta variants (4 ug/ml per peptide). The spots were developed based on the manufacturer’s instructions. In both animal models, PMA (50 ng/mL) and ionomycin (1 µM) were used as positive controls, and RPMI + 10% FBS with DMSO was used as the negative control. Spots were scanned and quantified using an S6 ImmunoSpot analyzer (CTL).

### FluoroSpot Assay for IFNγ, TNFα and IL2 to assess T cell activation

Precoated 96-well mouse IFNγ/TNFα/IL2 FluoroSpot plates (MabTech, cat# FSP-414245) were blocked with RPMI + 10% FBS for 2 hours at room temperature. 0.1×10^6^ mouse splenocytes were plated into each well and stimulated overnight with RBD- or OVA-derived OLPs (2 µg/ml per peptide). FluoroSpot assays were performed as instructed by the manufacturer.

### Multiplexed proteomics (Luminex) to assess T cell activation

To determine cytokine production from T cells, mice were immunized twice, and spleens were restimulated overnight with OLPs 7 days after booster dose. Cell culture supernatant was diluted 1:2 (TNFα, IL2, GM-CSF) or 1:20 (IFNγ, GzmB) and assayed using the MCD8MAG-48K kit (EMDMillipore). The assay was performed as instructed by the manufacturer.

To determine the cytokine/chemokine content of mouse LNs, animals were vaccinated once, and inguinal LNs were collected 6-48 hours post immunization. For IFNAR-1 blockade, an antagonistic mAb (clone: MAR1-5A3, BioXcell, cat# BE0241) or an isotype control (clone: MOPC-21, BioXcell, cat# BE0083) were injected intraperitoneally 24 hours prior to immunization. Protein Extraction Buffer (Invitrogen, cat# EPX-9999-000) containing Mini protease inhibitor cocktail (Roche, cat# 53945000) and HALT phosphatase inhibitors (Thermo Fisher, cat# 78442) was added to the intact LNs prior to homogenization using a TissueLyser II (Qiagen). Centrifugation-cleared lysates were analyzed with Luminex Cytokine and Chemokine kits (EMDMillipore, cat# MCYTOMAG-70K and MECY2MAG-73K).

### Enzyme linked immunosorbent assay (ELISA) for antibody titers

ELISAs were performed to determine antigen-specific antibody titers in serum. For murine studies, ELISA plates were coated with OVA (200 ng/well; InvivoGen; cat# vac-pova) or RBD antigen (200 ng/well; GenScript; cat# Z03483) overnight at 4 °C. Plates were pre-blocked with 2% bovine serum albumin for 2 h at room temperature (RT). Serially diluted mouse sera were transferred to the ELISA plates and incubated for 2 h at RT. Plates were washed four times with washing buffer (BioLegend; cat# 4211601) and then incubated for 1 h at RT with horseradish peroxidase (HRP)-conjugated anti-mouse secondary antibodies (1:2000 dilution) against either pan-IgG (Fcγ) (Jackson ImmunoResearch, cat# 315-035-046), IgG1 (Jackson ImmunoResearch, cat# 115-035-205), or IgG2c (Jackson ImmunoResearch, cat# 115-035-208). Plates were again washed four times with washing buffer, after which the plates were developed with 3,3′,5,5′-tetramethytlbenzidine for 10 min at RT, and the reaction was stopped with 1 N sulfuric acid. The absorbance at 450 nm was measured by an ELISA plate reader. Titers were determined at an absorbance cutoff of 0.5 OD (optical density).

For NHP assays, ELISA plates were coated with 100 ng/well of RBD WH-01, Beta, Delta, or Omicron antigen. The detection antibody used was horseradish peroxidase (HRP)–conjugated goat anti-human IgG (H+L) (ThermoFisher, cat# SA5-10283) at a 1:2000 dilution. The WHO International Standard for anti-SARS-CoV-2 immunoglobulin (anti-RBD IgG High 20/150, NIBSC) was used for reference values. Serum titers were determined at an absorbance cutoff of 0.5 OD and converted into binding antibody units/mL (BAU/mL) using the WHO standard.

### SARS-CoV-2 pseudovirus neutralization assay for NHP sera

The SARS-CoV-2 pseudovirus assay was performed by Genecopoeia as previously described^26^. SARS-CoV-2 Spike-pseudotype lentiviruses from the Washington 1 (WA1) D614G, Delta and Omicron variants were used. Briefly, a HEK293T cell line overexpressing ACE2 and TMPRSS2 was seeded at a density of 1.2×10^4^ cells/well overnight. 3-fold serial dilutions of heat inactivated serum samples were prepared and mixed with 50 µL of pseudovirus. The mixture was incubated at 37°C for 1 hour before adding to the HEK-293T-hACE2 cells. After 72 hours, the cells were lysed, and firefly luciferase activity was determined. SARS-CoV-2 neutralization titers were defined as the sample dilution at which a 50% reduction in RLU was observed relative to the average of the virus control wells. Convalescent plasma samples from patients who had recovered from SARS-CoV-2 infection (COVID-19) within 4 weeks of sample collection were obtained from US Biolab (Rockville, MD) and ALLCELLS (Alameda, CA). All samples were received and stored frozen at −80°C until analysis.

### Gene transcript analysis by Nanostring

For mouse studies, inguinal LNs were harvested from immunized C57BL/6J mice at the indicated time points, processed into single cell suspensions, and lysed with RLT buffer (Qiagen, cat# 79216). Transcriptional profiles of immune signaling were generated using the nCounter Mouse Immunology Panel of 561 mouse immune response genes (NanoString Technologies, cat# XT-CSO-MIM1-12). For NHP studies, LN FNA were assessed using the nCounter NHP Immunology Panel of 770 macaque immune response genes (NanoString Technologies, cat# XT-CSO-NHPIM1-12). Transcriptional responses were assessed with nSolver software v4.0 (NanoString Technologies) and differential gene expression analysis was carried out using ROSALIND software (Rosalind, Inc).

### Biodistribution analysis by IVIS

C57BL/6J mice were immunized once with 5 μg WH-01 Spike RBD protein and 5 nmol Cy5-labelled DNA adjuvant. ‘No Cy5’-control animals were administered antigen plus unlabeled AMP-DNA. 6 h, and 1, 3, 5, and 7 days post injection, inguinal LNs were harvested and imaged *ex vivo* using the IVIS Spectrum (PerkinElmer, Waltham, MA, USA).

### Statistics

For comparing two experimental groups, two-tailed t-test analysis was utilized when normal distribution and homogeneity of variance, determined by Levene’s Test, were established. Where these assumptions did not apply, the Mann-Whitney test was used instead. For comparison of multiple groups, ordinary one-way ANOVA followed by Tukey’s, or Šidák’s post-hoc analysis, as indicated, was used to compare experimental groups. NanoString statistical analysis was performed using Rosalind software.

## Data Availability

The datasets supporting the findings presented in this study are available from the corresponding author upon reasonable request. Nanostring data is available in the Gene Expression Omnibus (GEO) under accession numbers GSE280859 and GSE280856 for the mouse and NHP data sets, respectively. All requests for data will be promptly reviewed by Elicio Therapeutics, Inc. to verify whether the request is subject to any intellectual property obligations or otherwise. Any data that can be shared will be released via a written agreement.

## ACKNOWLEDGEMENTS

We thank Crystal Carter, Jane Fontenot, Francois Villinger at New Iberia Research Center for the execution of the NHP studies; the NIH Tetramer Core Facility at Emory University for supplying the SARS-CoV-2 RBD tetramer; and Darrell J. Irvine at the Massachusetts Institute of Technology/Scripps Research Institute for helpful advice and discussion. We acknowledge funding support from Elicio Therapeutics.

## AUTHOR INFORMATION

### Contributions

**M.P.S.** conceived and led the study, designed and performed experiments, analyzed and interpreted data, and wrote the manuscript. **L.M.S.** performed immunogenicity and transcriptome experiments and analyzed data. **W.Z** designed and performed murine knock-out experiments and analyzed data. **E.P.** performed immunogenicity and transcriptome experiments. **A.J.** conceived, designed and performed proteomics and NHP serum cytokine experiments, analyzed and interpreted data. **X.C.-P.** and **M.M.J.** performed immunogenicity experiments and analyzed data. **L.K.M.** conceived and designed NHP experiments, interpreted data. **C.M.H.** advised on study design. **K.A.F.** conceived and designed murine knock-out experiments, analyzed and interpreted data. **P.C.D.** conceived and led the study, designed experiments, analyzed and interpreted data. All authors reviewed and approved the final manuscript.

## SUPPLEMENTARY FIGURES

**Fig S1:**
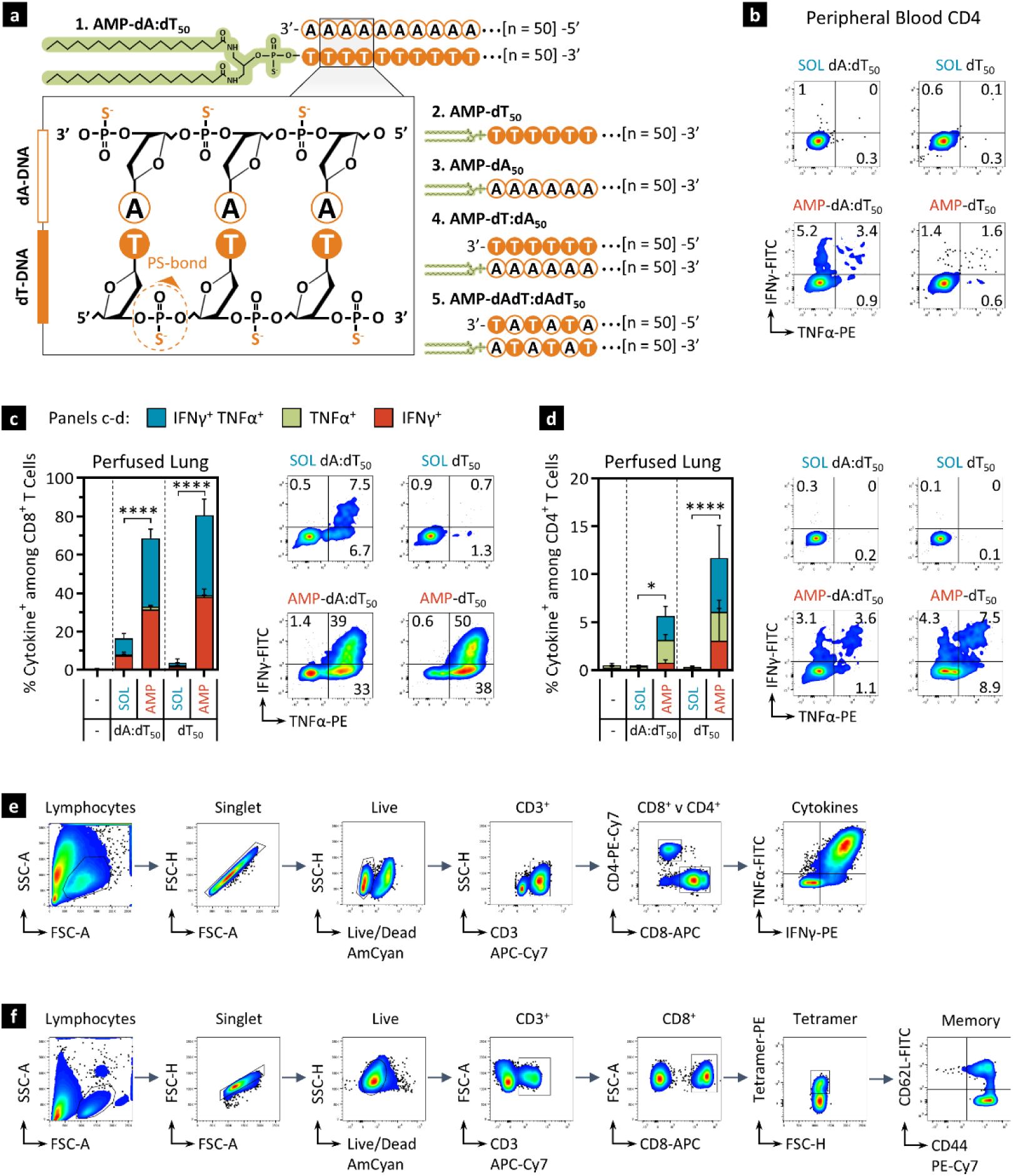
AMP-DNA is a potent adjuvant for inducing responses to protein subunit antigens – Additional data for Figure 1. **a)** Schematic drawings depicting the chemical structure of the AMP-DNA adjuvants. The ssDNA variants, dT_50_ and dA_50_, consist of 50 phosphorothioate (PS)-linked deoxythymidines and deoxyadenosines, respectively. When annealed to each other the resulting dsDNA forms dA:dT_50_. The diacyl lipid (AMP-tail) is conjugated to the 5’-end of each ssDNA variant, or to the dT strand in dA:dT_50_. The dT:dA_50_ molecule is AMP-conjugated on the dA strand. The dAdT:dAdT_50_ molecule consists of alternating deoxyadenosine and deoxythymidine residues. **b-d)** C57BL/6J mice (n = 5) were immunized twice with 5 μg OVA protein and 5 nmol SOL or AMP-DNA adjuvants as in Fig. 1. Cells were assayed 7 days post booster dose. b) Example flow cytometric scatter plots of IFNγ and TNFα production by CD4^+^ T cells in peripheral blood (see Fig. 1i). b,c) flow cytometric analysis of IFNγ and TNFα production in CD8^+^ and CD4^+^ T cells from perfused lung tissue. **e-f)** Gating strategy for intracellular cytokine staining of T cells re-stimulated with peptides overnight (e), and gating strategy for tetramer^+^ CD8^+^ T cells and memory phenotypes of freshly processed PBMCs (f).

**Fig S2:**
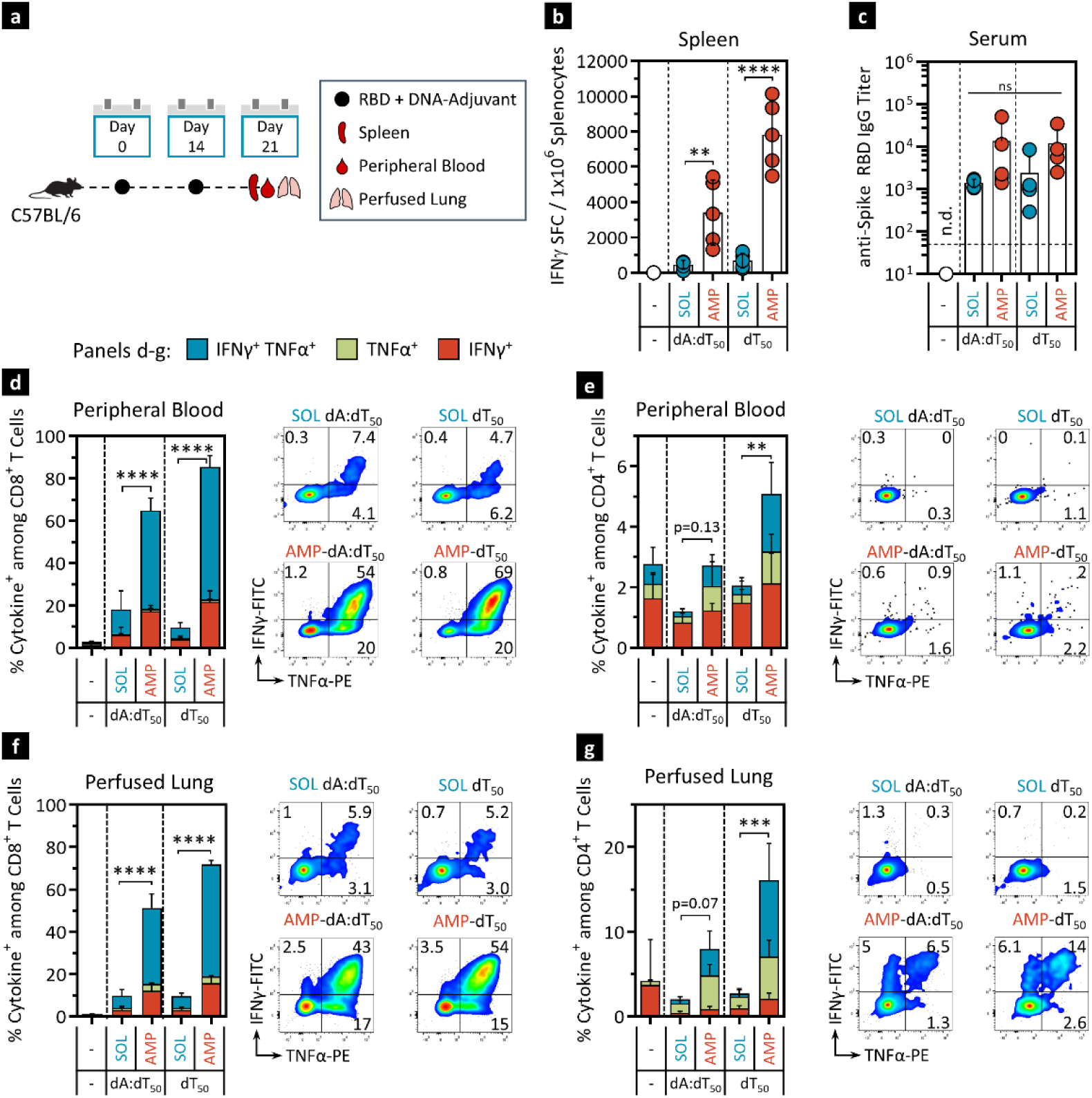
AMP-DNA is a potent adjuvant for inducing T cell and Ab responses to the RBD protein subunit antigen: **a)** Schema showing animal dosing and experimental schedule. C57BL/6J mice (n = 5) were immunized twice with 5 μg WH-01 RBD protein and 5 nmol SOL or AMP-DNA adjuvants. Cells were assayed 7 days post booster dose. **b)** Splenocytes were restimulated with RBD OLPs overnight and assayed for IFNγ production by ELISpot. Shown is the frequency of IFNγ SFCs per 10^6^ splenocytes. **c)** Serum anti-RBD IgG titers were determined against WH-01 RBD protein. **d-g)** Flow cytometric analysis of cytokine production by CD8^+^ (d, f) and CD4^+^ T cells (e, g) in peripheral blood and perfused lung tissue. Shown are percentages of cytokine^+^ cells among CD8^+^ or CD4^+^ T cells and representative flow cytometry dot plots of TNFα and IFNγ positive T cells. Mock vaccines contained vehicle only. Values depicted are means±SD. *p < 0.05, **p < 0.01, ***p < 0.001, ****p < 0.0001 by one-way ANOVA followed by Tukey’s post-hoc analysis. n.d., not detected.

**Fig S3:**
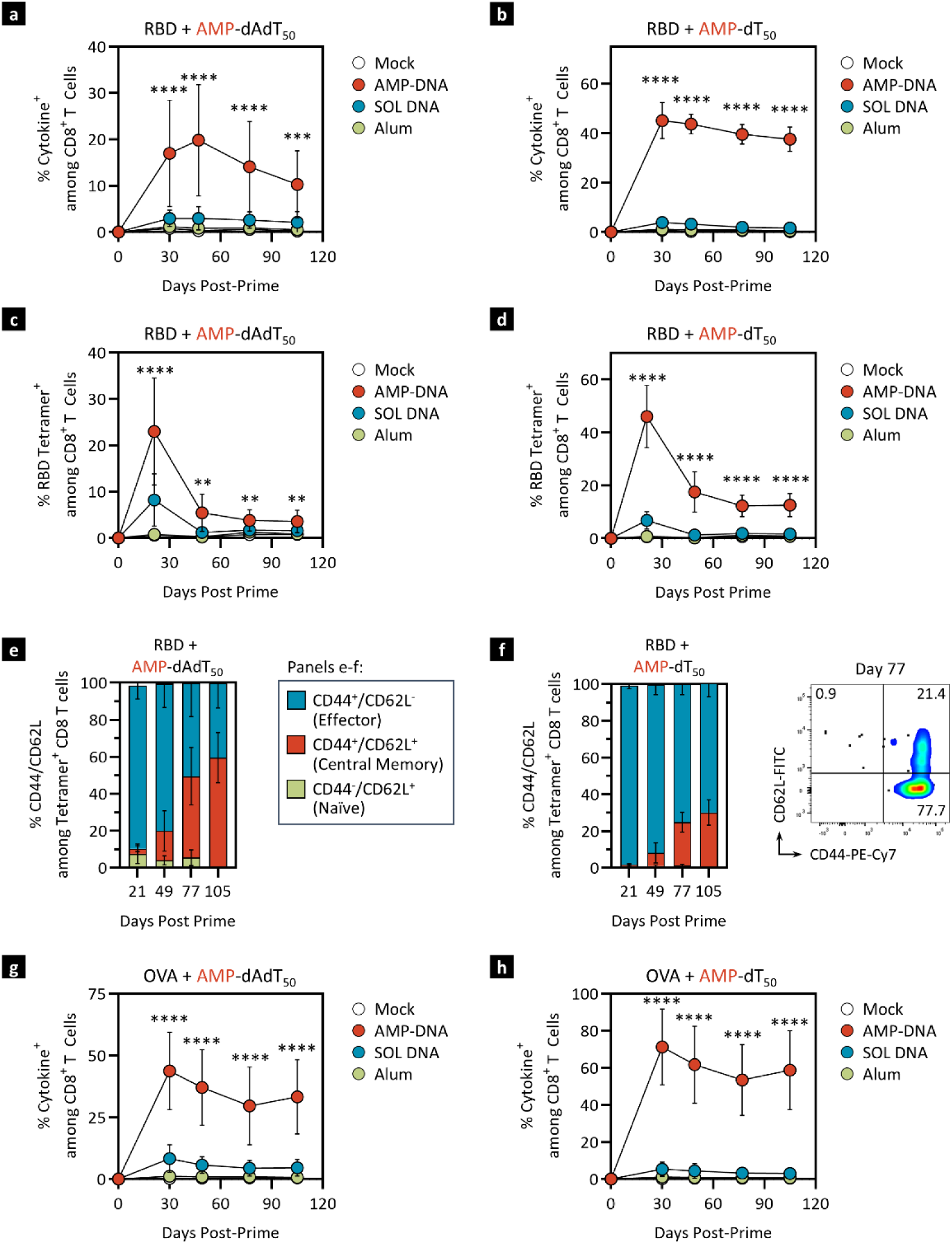
Long-term T cell response to AMP-DNA adjuvants. **a-f)** C57BL/6J mice (n = 10) were immunized twice with 5 μg WH-01 RBD protein adjuvanted with either 100 μg alum or 5 nmol dAdT_50_ (a,c,e), or dT_50_ (b,d,f). Mock vaccines contained vehicle only. Peripheral blood samples were collected, processed and stimulated with RBD OLPs overnight at the indicated time points. a,b) Flow cytometric analysis of cytokine production by CD8^+^ T cells in peripheral blood. Shown are percentages of cytokine^+^ cells among CD8^+^ T cells. c,d) Peripheral blood CD8^+^ T cells were assayed for RBD reactive TCRs using a VNFNFNGL-specific tetramer. Shown are Tetramer^+^ cells among CD8 T cells. e,f) Tetramer^+^ CD8^+^ T cells from peripheral blood were stained for CD44 and CD62L to determine memory phenotype. **g-h)** C57BL/6J mice (n = 10) were immunized twice with 5 μg OVA protein adjuvanted with either 100 μg alum or 5 nmol dAdT_50_ (g), or dT_50_ (h). Mock vaccines contained vehicle only. Peripheral blood samples were collected, processed and stimulated with OVA OLPs overnight at the indicated time points. Statistics in a-d) and g-h) compare AMP- to SOL DNA treatment groups. Values depicted are means±SD. *p < 0.05, **p < 0.01, ***p < 0.001, ****p < 0.0001 by one-way ANOVA followed by Tukey’s post-hoc analysis.

**Fig S4:**
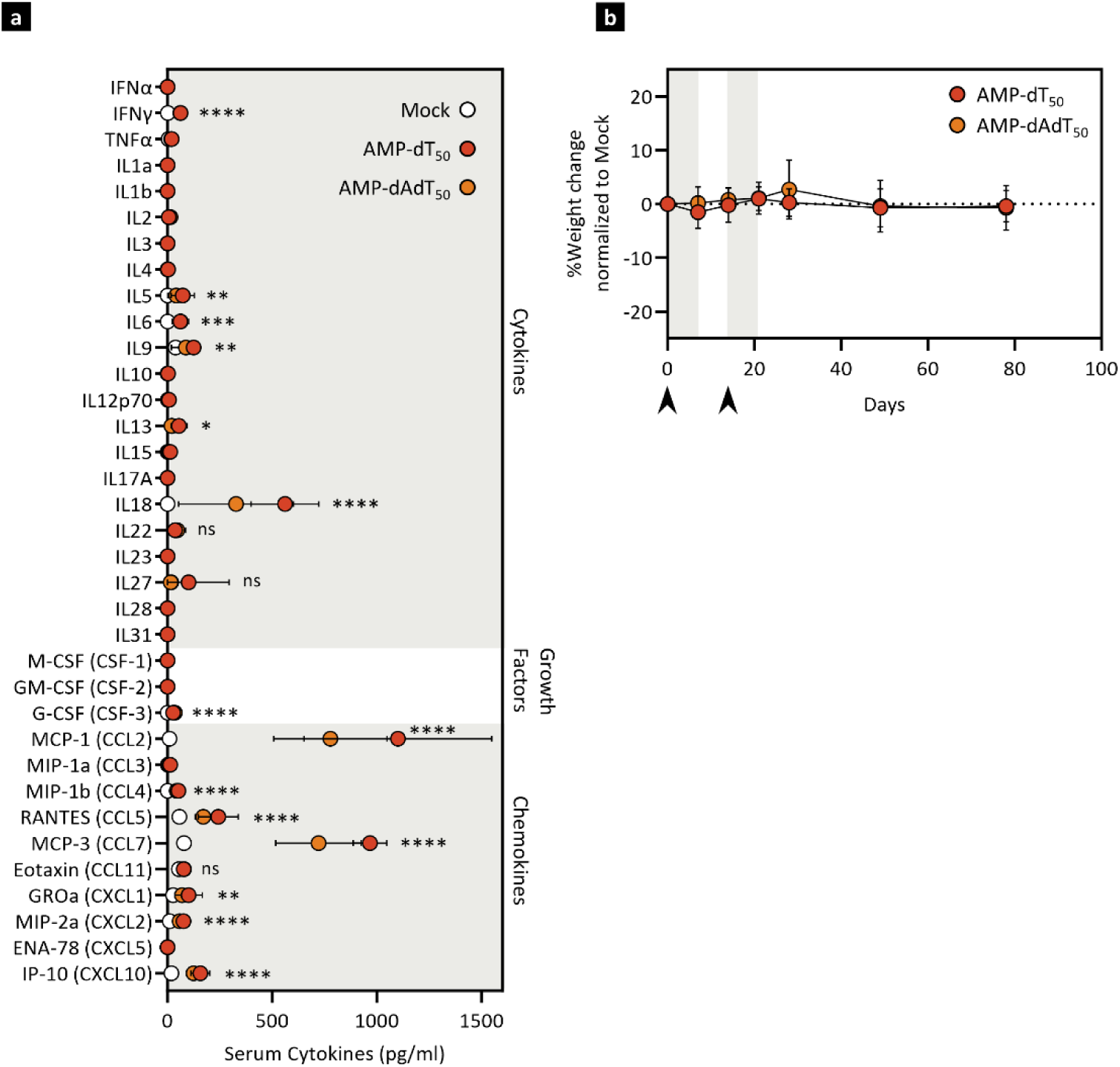
No adverse serum cytokine levels or weight changes are observed upon administration of AMP-DNA adjuvants in mice. C57BL/6J mice (n = 10) were immunized once with 5 μg OVA protein and 5 nmol AMP-dT_50_ or AMP-dAdT_50_. Mock vaccines contained vehicle only. **a)** Serum was collected 24 hours after immunization and analyzed by Luminex for 35 inflammatory cytokines, chemokines and growth factors. Values depicted are means±SD. *p < 0.05, **p < 0.01, ***p < 0.001, ****p < 0.0001 by one-way ANOVA followed by Tukey’s post-hoc analysis comparing AMP-DNA treated groups to Mock. **b)** Animal weight was recorded at indicated time points and is depicted as mock-subtracted percent change from baseline (day 0). Arrows indicate immunization timepoints. Gray shading in b) indicates one week post each immunization.

**Fig S5:**
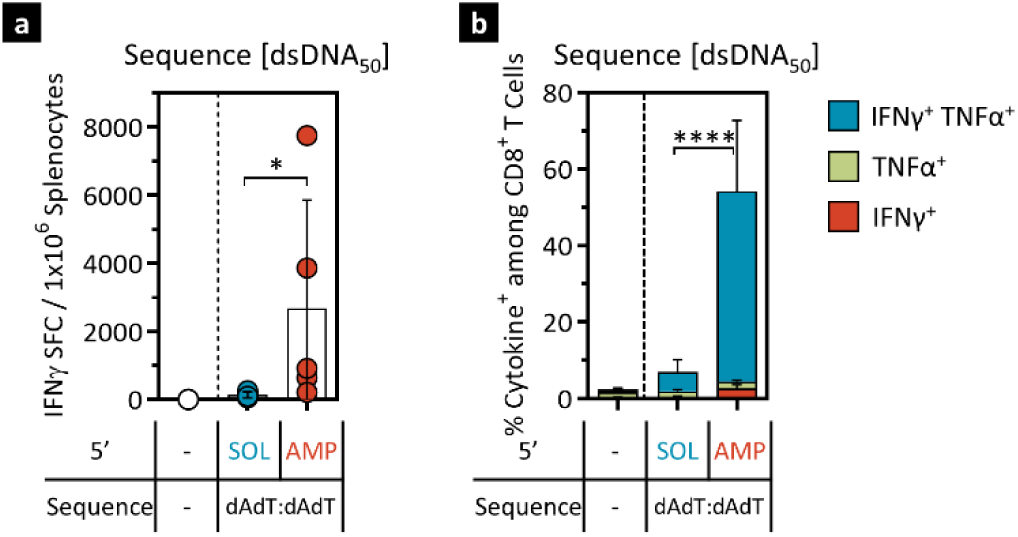
AMP-dAdT:dAdT rescues AMP-dA responses – Additional data for Figure 2. C57BL/6J mice (n = 5) were immunized as described in Fig 1c. Immunizations contained 5 μg WH-01 RBD protein and 5 nmol SOL or AMP-dAdT:dAdT adjuvant. Cells were assayed 7 days post booster dose. **a)** Splenocytes were restimulated with RBD OLPs overnight and assayed for IFNγ production by ELISpot. Shown is the frequency of IFNγ SFCs per 10^6^ splenocytes. **b)** Flow cytometric analysis of cytokine production by CD8^+^ T cells in peripheral blood. Shown are percentages of cytokine^+^ cells among CD8^+^ T cells. Mock vaccines contained antigen only. Values depicted are means±SD. *p < 0.05, **p < 0.01, ***p < 0.001, ****p < 0.0001 by one-way ANOVA followed by Tukey’s post-hoc analysis.

**Fig S6:**
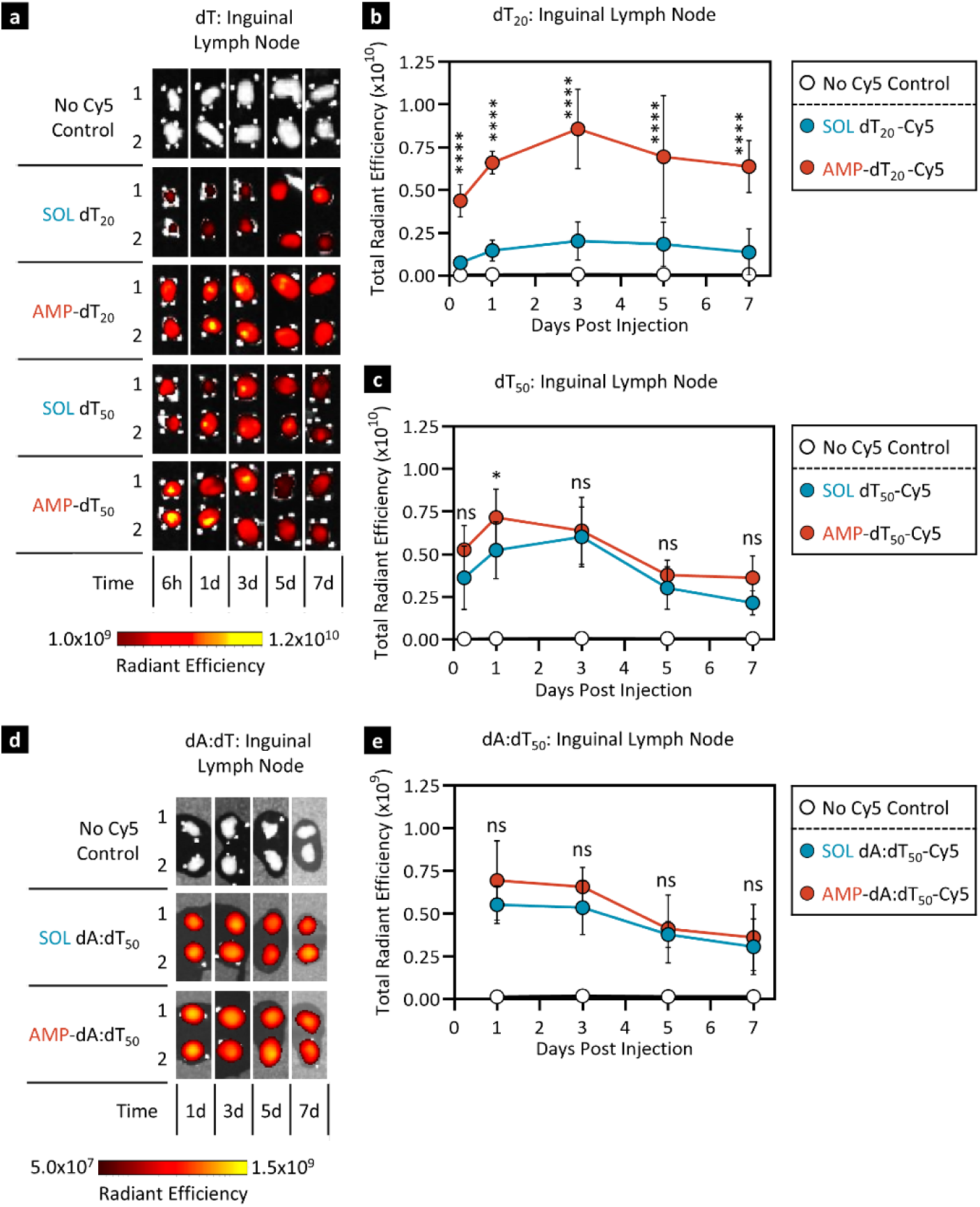
AMP-mediated lymph node delivery of DNA is length dependent. C57BL/6J mice (n = 5 per time point) were immunized once with 5 μg WH-01 RBD protein and 5 nmol Cy5 fluorophore-labelled DNA-adjuvant. Lymph nodes were harvested and assayed via IVIS at the indicated time points post immunization. ‘No Cy5’-controls contained antigen and unlabeled AMP-dT_50_. **a)** Representative IVIS images of extracted inguinal lymph nodes from mice treated with SOL or AMP-dT length variants (dT_20_ and dT_50_). **b, c)** Quantification of total radiant efficiency from panel a) for dT_20_ (b) and dT_50_ (c). **d)** Representative IVIS images of extracted inguinal lymph nodes from mice treated with SOL or AMP-dAdT_50_. **e)** Quantification of total radiant efficiency from panel d). Values depicted are means±SD. *p < 0.05, **p < 0.01, ***p < 0.001, ****p < 0.0001 by one-way ANOVA followed by Šidák’s post-hoc analysis. Statistical analyses compare SOL DNA-adjuvants to their AMP-modified counterparts.

**Fig S7:**
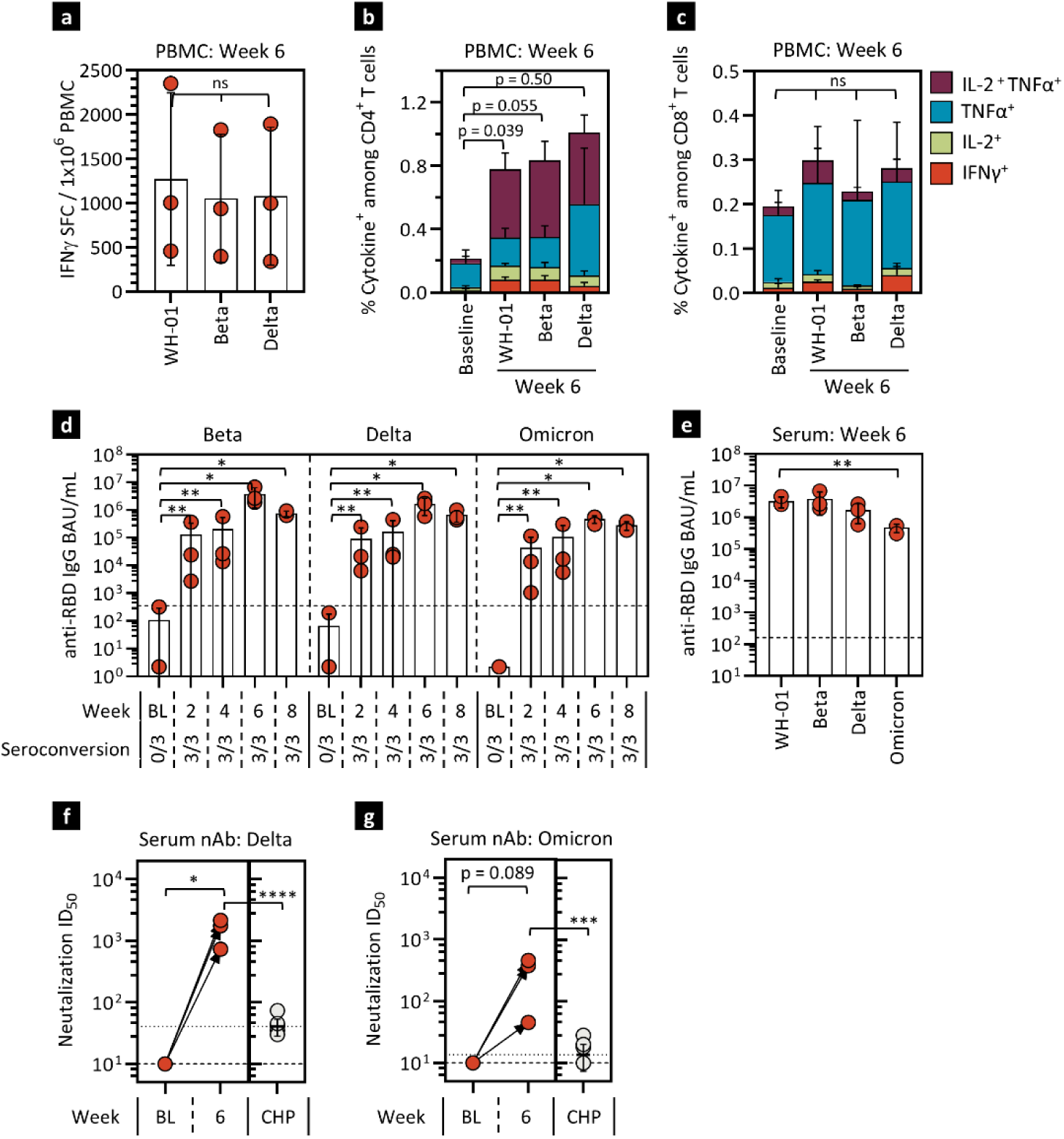
AMP-dT generates cross-protective cellular and humoral responses to SARS-CoV-2 variants of concern in NHP. Cellular and humoral responses to RBD and AMP-DNA adjuvant were assessed in rhesus macaques as in Fig. 4. Three animals were immunized at week 0 and 4 with 140 µg WH-01 RBD protein admixed with 5 mg of AMP-dT_50_. **a-c)** T cell cross-reactive responses were compared upon overnight restimulation with RBD OLPs to variants of concern (VOC) at week 6 by a) IFNγ ELISpot analysis [shown are IFNγ SFCs per 1 x 10^6^ PBMCs], and b-c) flow cytometric analysis of cytokine production by CD4^+^ and CD8^+^ T cells [shown are frequencies of IFNγ, TNFα and IL2 cytokine production]. **d)** RBD-specific serum IgG binding antibody units (BAU) were assessed for VOCs at each sample collection time point and **e)** compared among VOCs at week 6. **f, g)** Neutralizing antibody responses were assessed through pseudovirus inhibition at week 6. Shown are ID_50_ values for Delta and Omicron variants in comparison to convalescent human plasma (CHP). The dashed lines represent the lower limit of detection discriminating between samples positive or negative for seroconversion and dotted lines represent the mean value observed for the human plasma comparators where appropriate. Values depicted are mean±SD. *\* p < 0.05; ** p < 0.01; *** p < 0.001; **** p < 0.0001* by paired t test.

**Fig S8:**
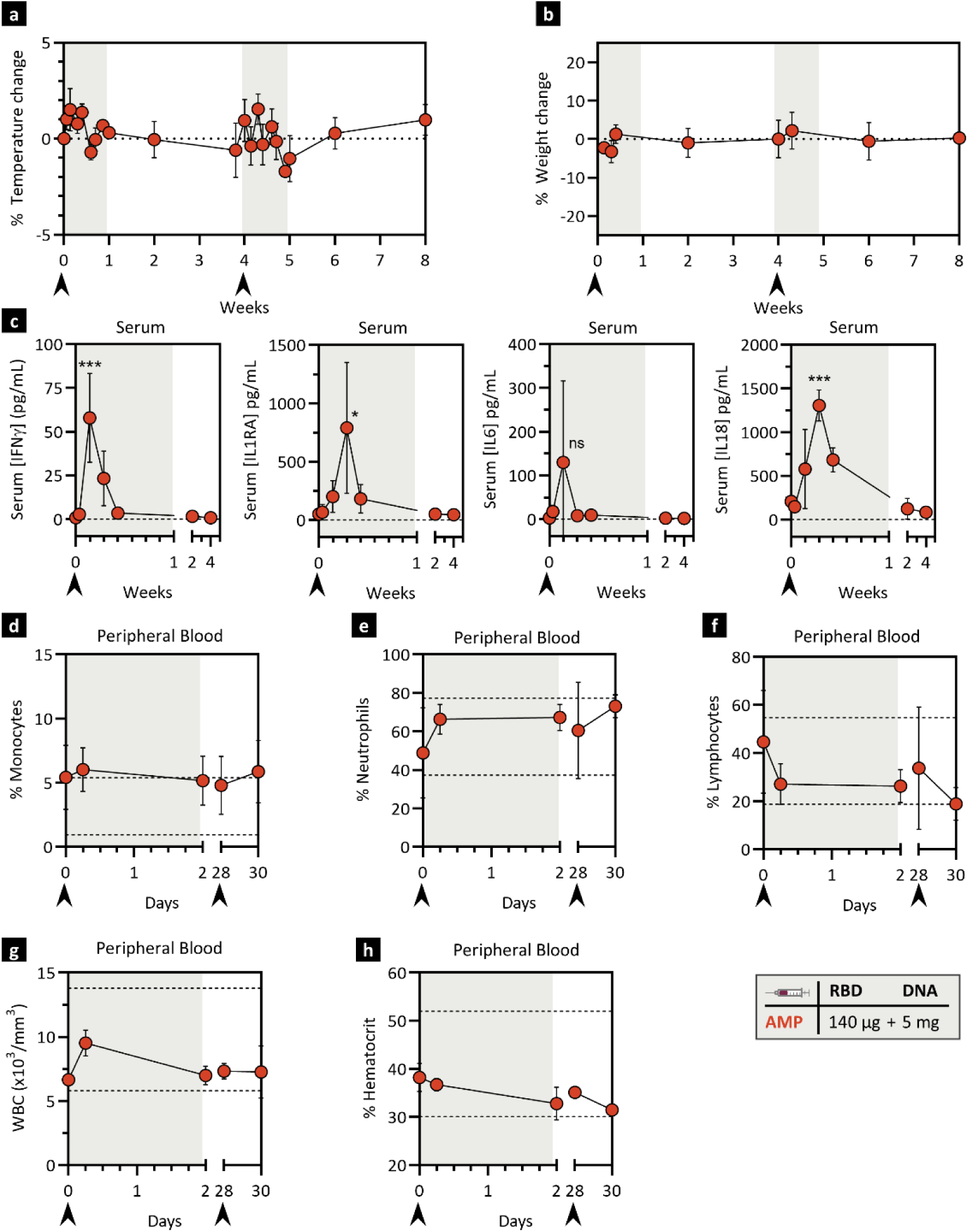
AMP-dT induces transient serum cytokine elevation in NHPs. Rhesus macaques (n=3) were immunized at week 0 and 4 with 140 µg WH-01 RBD protein admixed with 5 mg of AMP-dT_50_. **a-b**) Clinical observations were recorded. Shown are percent change for temperature (a) and weight (b) from baseline. **c**) Sera were collected at multiple timepoints for cytokine assessment by Luminex. The dashed lines in a-c indicate no change from baseline. Significance is based on baseline value. **d-h**) Sera were collected at multiple timepoints for assessment in a hematology complete blood count panel. Shown are % monocytes (d), % neutrophils (e), % lymphocytes (f), white blood cells (WBC) (g) and % hematocrit (h). Dashed lines indicate reference value ranges. Arrows under each graph indicate immunization timepoints. Gray shading indicates one week post each immunization.

**Fig S9:**
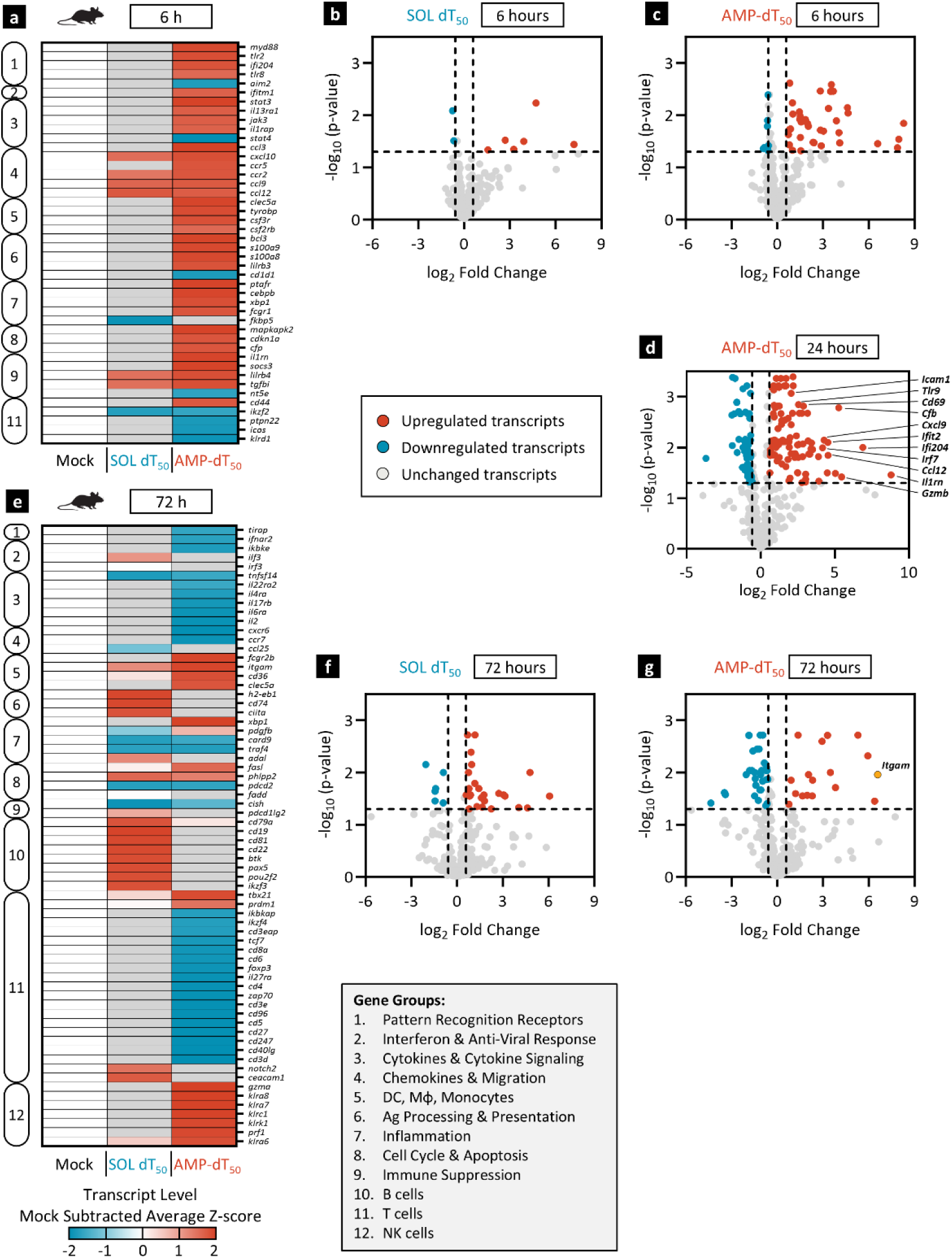
AMP-DNA promotes comprehensive inflammatory transcriptional reprogramming in draining lymph nodes of mice. Mouse LN samples were collected and analyzed as in Fig. 5. **a,e)** Heatmap representation of whole LN mRNA analyzed by NanoString nCounter Mouse Immunology Panel 6 hours and 72 hours after immunization, respectively. Shown are mock-subtracted, average Z-scores. Mock vaccines contained vehicle only. Shown are transcripts out of 567 total genes that are significantly changed in at least one treatment group (p < 0.05 as determined with Rosalind software) and that are either ≥1.5-fold upregulated (red) or downregulated (blue) compared to Mock treatment. Insignificant values (p ≥ 0.05) are shown in gray. Genes were clustered into 12 groups (box insert) using Gene Ontology and UniProt databases. **b-d,f-g)** Volcano plot representation of log-transformed transcript values: b) SOL dT_50_ at 6 hours, c) AMP-dT_50_ at 6 hours, d) AMP-dT_50_ at 24 hours (same data as Fig. 5c with additional gene annotations), f) SOL dT_50_ at 72 hours, g) AMP-dT_50_ at 72 hours (red values are significantly upregulated and blue values are significantly downregulated gene transcripts). Dashed horizontal line represents significance threshold (p = 0.05); vertical dashed lines represent fold-change limits of ±1.5.

**Fig S10:**
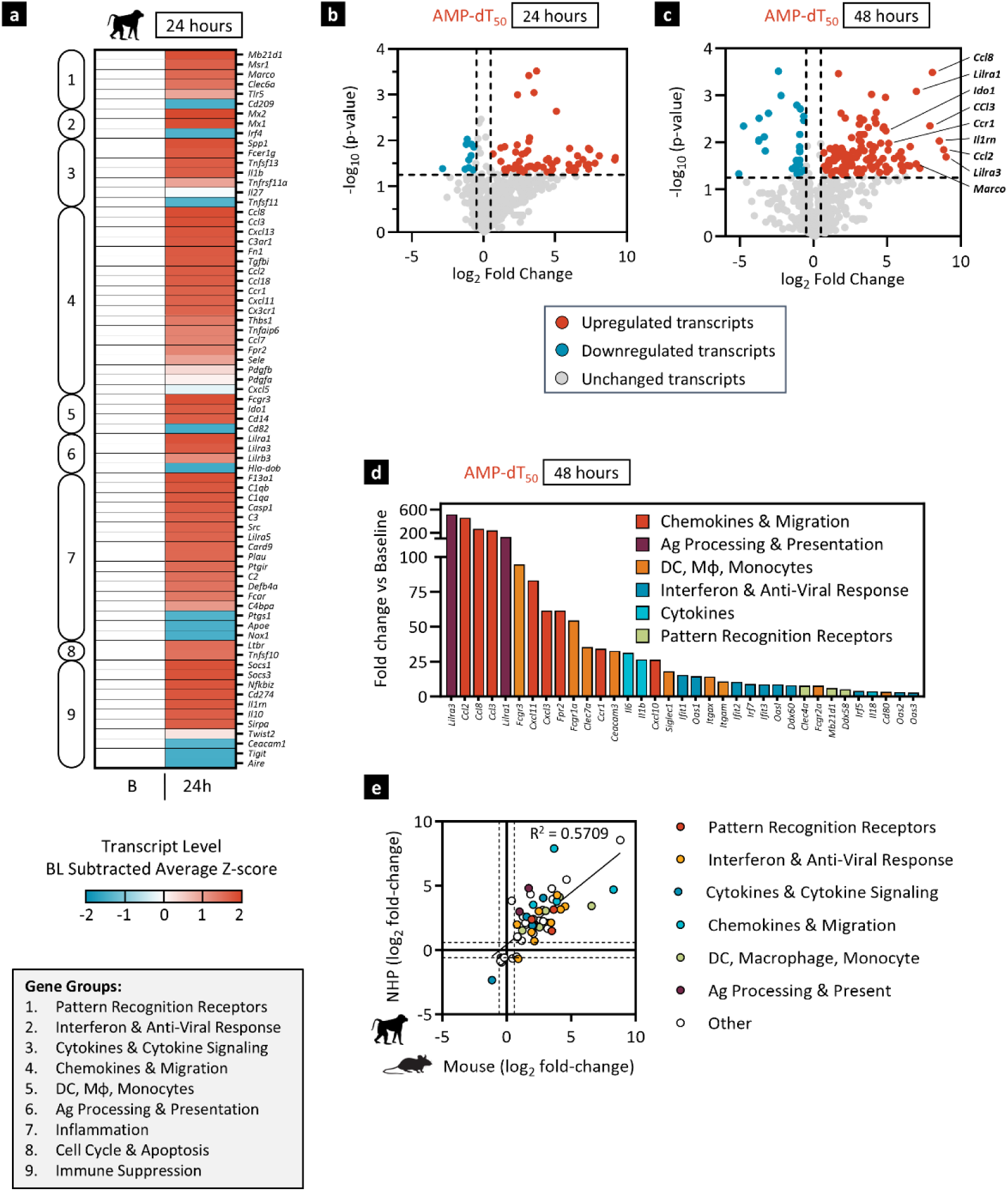
AMP-DNA promotes comprehensive inflammatory transcriptional reprogramming in draining lymph nodes of NHP. NHP LN FNA samples were collected and analyzed as in Fig. 5. **a)** Heatmap representation of LN FNA mRNA analyzed by NanoString nCounter NHP Immunology Panel 24 hours after immunization. Shown are baseline-subtracted, average Z-scores of transcripts that are significantly changed (p < 0.05 as determined with Rosalind software) and are either ≥1.5-fold upregulated (red) or downregulated (blue) compared to baseline. Genes were clustered into 12 groups (box insert) using Gene Ontology and UniProt databases. **b-c)** Volcano plot representation of log-transformed transcript values from AMP-dT_50_ immunized NHP at b) 24 hours and c) 48 hours after immunization (same data as Fig. 5e with additional gene annotations). Red values are significantly upregulated and blue values are significantly downregulated gene transcripts that changed by a minimum of 1.5-fold. Dashed horizontal line represents significance threshold (p = 0.05); vertical dashed lines represent fold-change limits of ±1.5. **d)** Fold-change bar graphs for selected NHP gene transcripts at 48 hours post boost. **e)** Scatter plot of genes that are significantly affected by AMP-dT_50_ immunization in both animal models at any time point and changed at least 1.5-fold in either model. Dashed lines represent fold-change limits of ±1.5.

**Fig S11:**
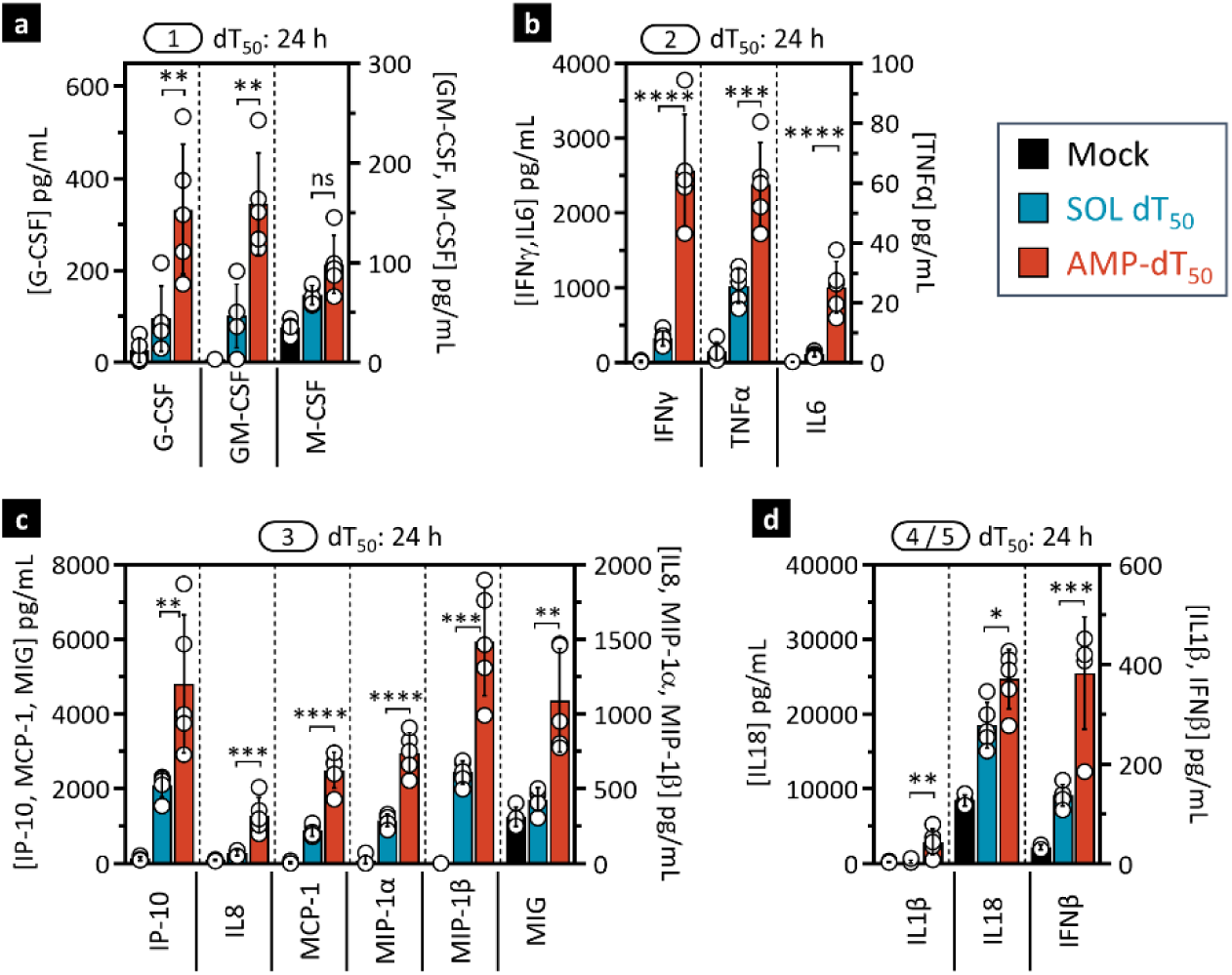
AMP-dT_50_ potently induces inflammatory proteomic milieu in draining lymph nodes. **a-d)** As in Fig. 6a, C57BL/6J mice (n = 5) were immunized once with 5 μg WH-01 RBD protein and 5 nmol SOL or AMP-conjugated dT_50_. Lymph nodes were collected and processed for protein extraction and analyzed by Luminex at 24 hours post immunization. Shown are concentrations of measured analytes (from Fig. 6a) separated by functional groups: (1) growth factors, (2) inflammatory cytokines, (3) chemokines, (4) inflammasome, and (5) IFN-I. Mock immunized animals received vehicle only. Values depicted are mean±SD. *p < 0.05; **p < 0.01; ***p < 0.001; ****p < 0.0001 by one-way ANOVA followed by Tukey’s post-hoc analysis.

**Fig S12:**
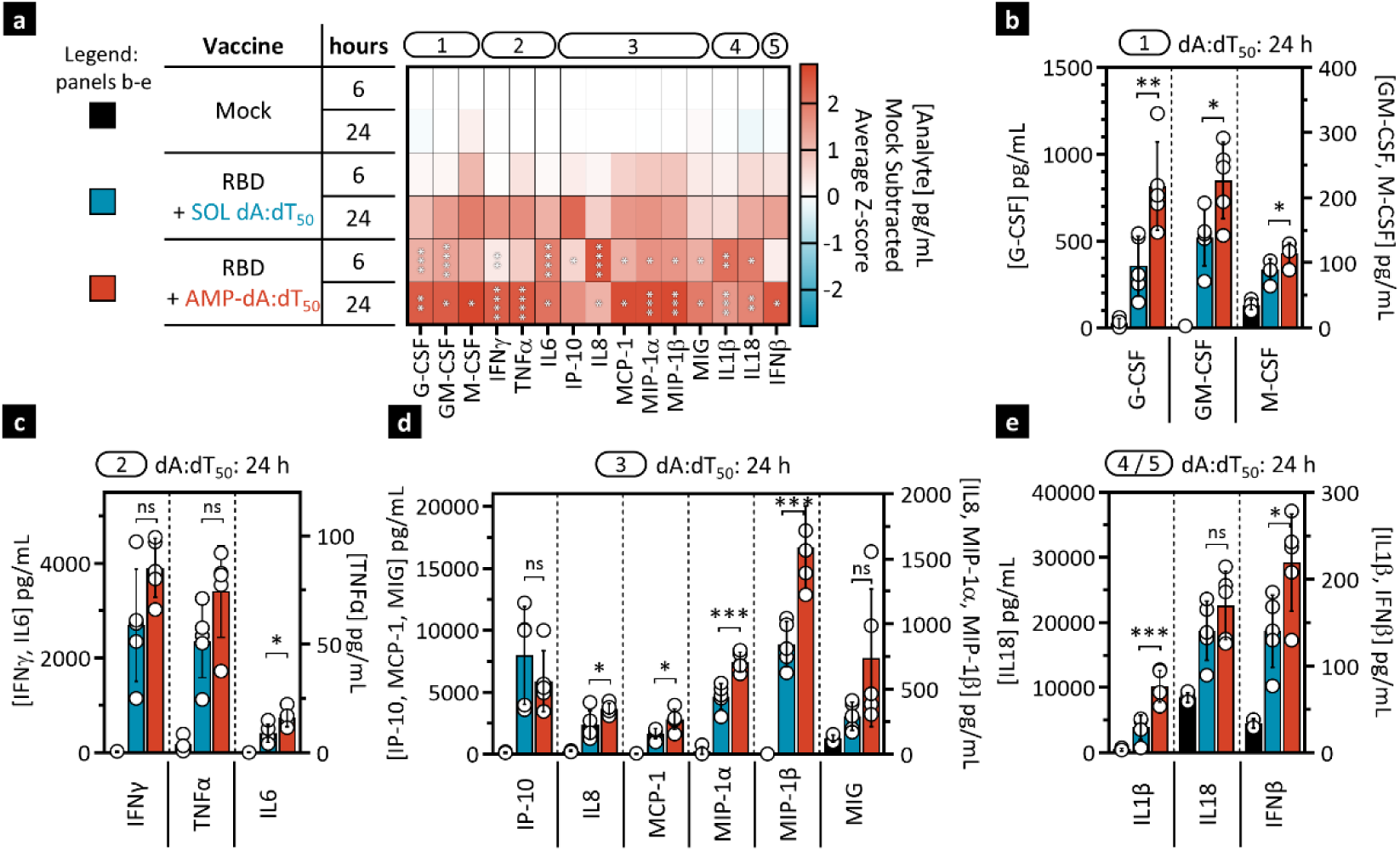
AMP-dAdT_50_ potently induces inflammatory proteomic milieu in the draining lymph nodes. C57BL/6J mice (n = 5) were immunized once with 5 μg WH-01 RBD protein and 5 nmol SOL or AMP-conjugated dAdT_50_. Lymph nodes were collected and processed for protein extraction and analyzed by Luminex at 6 and 24 hours post immunization. **a)** Heat map representation of mock subtracted Z-scores of analyte concentrations (pg/ml). Analytes are annotated by functional category as in Fig 6. **b-e)** Concentrations of measured analytes for functional groups 1 to 5 as indicated, at 24 hours post immunization. Mock immunized animals received vehicle only. Values depicted are mean±SD. *p < 0.05; **p < 0.01; ***p < 0.001; ****p < 0.0001 by one-way ANOVA followed by Tukey’s post-hoc analysis.

**Fig S13:**
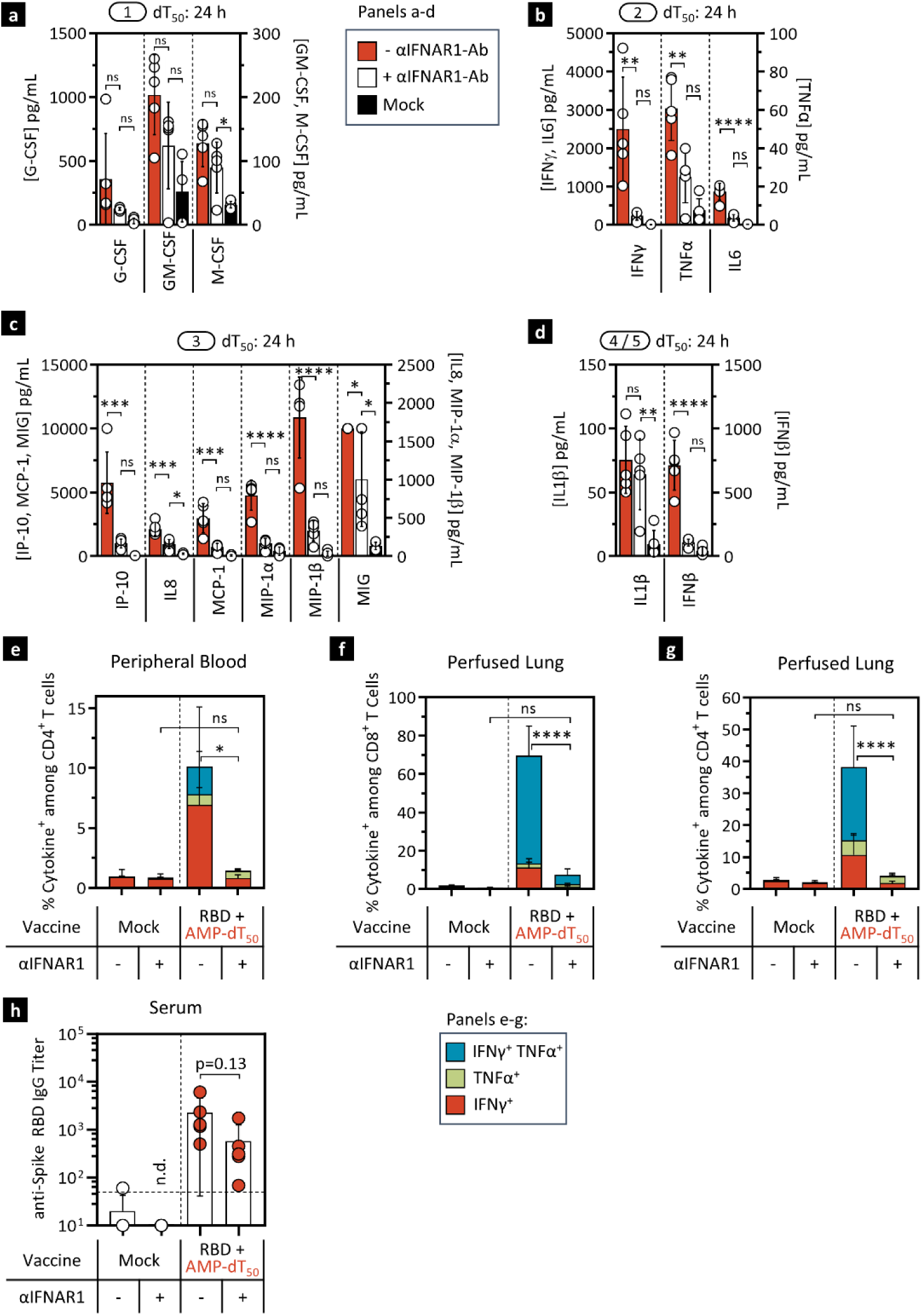
AMP-dT_50_-mediated immune activation in lymph nodes and immunogenicity is dependent on interferon signaling. **a-d)** As in Fig. 6b, C57BL/6J mice (n = 5) were dosed intraperitoneally with IFNAR1 blocking antibody or isotype control one day prior to immunization with one dose of 5 μg WH-01 RBD protein and 5 nmol AMP-dT_50_. Protein from lymph nodes was extracted 24 hours post immunization and analyzed by Luminex. Shown are concentrations of measured analytes (from Fig. 6b) separated by functional groups: (1) growth factors, (2) inflammatory cytokines, (3) chemokines, (4) inflammasome, and (5) IFN-I. **e-h)** As in Fig. 6c-d, C57BL/6J mice (n = 5) were immunized twice with 5 μg WH-01 RBD protein and 5 nmol AMP-dT_50_. Mice received IFNAR1 blocking antibody or isotype control intraperitoneally one day prior to each dose. Cells were assayed 7 days post booster dose. e-g) Shown are flow cytometric analyses of cytokine production by CD4^+^ T cells in peripheral blood (e), and CD8^+^ (f) and CD4^+^ (g) lung-resident T cells. h) Serum anti-RBD IgG titers were determined against WH-01 RBD protein. Mock immunized animals received vehicle only. Values depicted are mean±SD. *p < 0.05; **p < 0.01; ***p < 0.001; ****p < 0.0001 by one-way ANOVA followed by Tukey’s post-hoc analysis. n.d., not detected.

**Fig S14:**
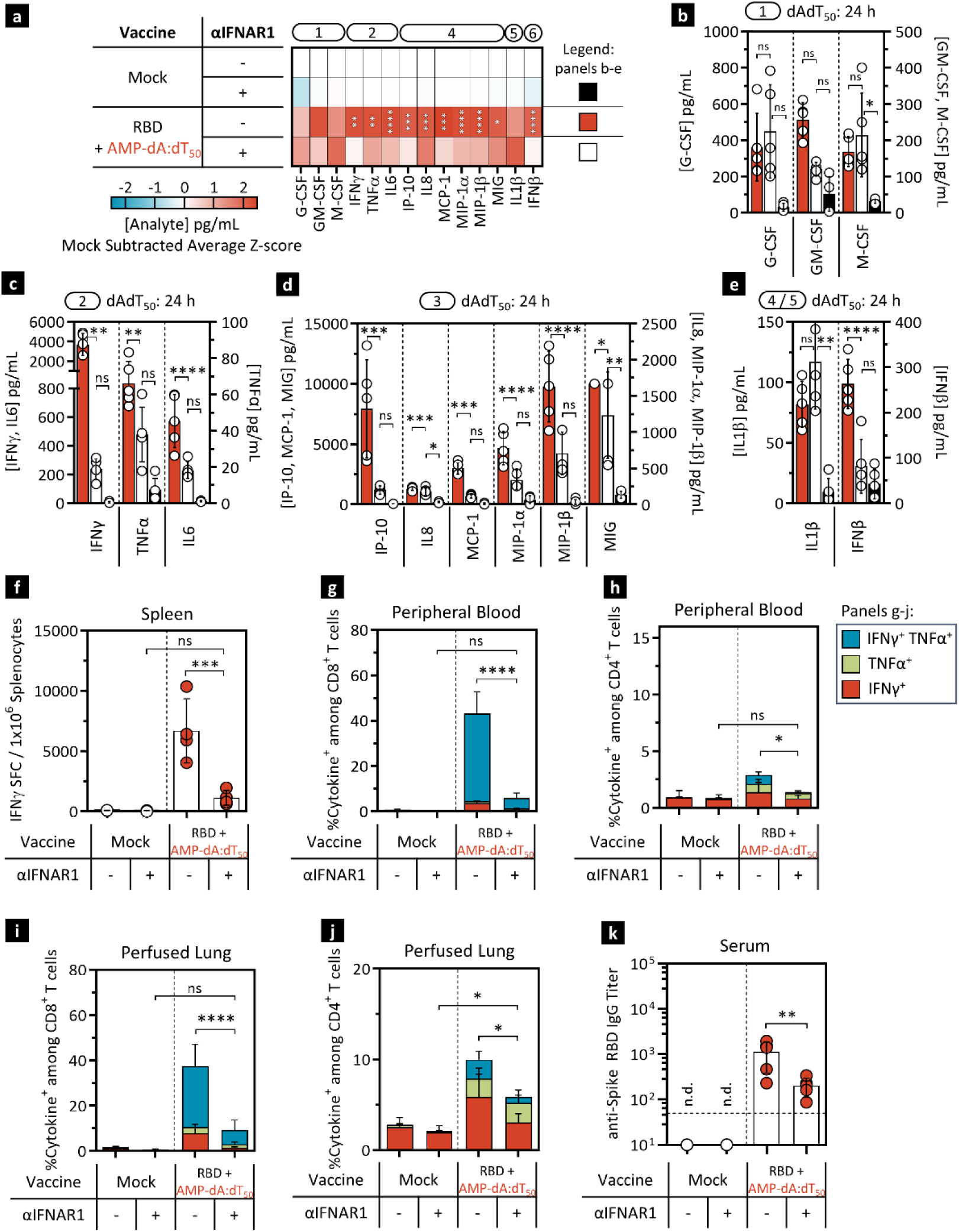
AMP-dAdT_50_-mediated immune activation in lymph nodes and immunogenicity is dependent on interferon signaling. **a-e)** C57BL/6J mice (n = 5) were dosed intraperitoneally with IFNAR1 blocking antibody or isotype control one day prior to immunization with one dose of 5 μg WH-01 RBD protein and 5 nmol AMP-dAdT_50_. Protein from lymph nodes was extracted 24 hours post immunization. a) Luminex assessed analyte concentrations were expressed in a heat map showing mock-subtracted Z-scores of analyte concentration (pg/ml). Functional group annotations as in Fig. 6b. b-e) Concentrations of measured analytes from panel a) for functional groups 1 to 5 as indicated. **f-k)** C57BL/6J mice (n = 5) were immunized twice with 5 μg WH-01 RBD protein and 5 nmol AMP-dAdT_50_. Mice received IFNAR1 blocking antibody or isotype control intraperitoneally one day prior to each dose. Cells were assayed 7 days post booster dose. f) Splenocytes were restimulated with RBD OLPs overnight and assayed for IFNγ production by ELISpot. Shown is the frequency of IFNγ SFCs per 10^6^ splenocytes. g, h) Flow cytometric analysis of cytokine production by CD8^+^ (g) and CD4^+^ T cells (h) in peripheral blood. k) Serum anti-RBD IgG titers were determined against WH-01 RBD protein. Mock immunized animals received vehicle only. Values depicted are mean±SD. *p < 0.05; **p < 0.01; ***p < 0.001; ****p < 0.0001 by one-way ANOVA followed by Tukey’s post-hoc analysis. n.d., not detected.

